# Tracking 3D forest density dynamics in a mixed temperate forest using occlusion-aware voxel transmittance and consistent multi-temporal UAV-LiDAR

**DOI:** 10.64898/2026.09.16.751970

**Authors:** Matthias Gassilloud, Barbara Koch, Anna Göritz

**Author notes:** Corresponding author: Email addresses (Matthias Gassilloud). (Barbara Koch), (Anna Göritz).

## Abstract

Forest structure and density dynamics drive ecosystem processes including carbon sequestration, microclimate regulation, and photosynthetic capacity. While LiDAR can capture these properties, high-frequency monitoring remains constrained by scarce multi-temporal datasets, inconsistent acquisitions, and occlusion biases confounding true structural change with methodological artifacts. This study investigates spatio-temporal forest structure dynamics through voxel transmittance in a European mixed temperate forest. We acquired an extensive UAV LiDAR dataset of 38 flights with identical sensor settings over more than two years. With a custom ray-tracing framework, we estimated voxel transmittance and mapped occlusion at 0.25 m resolution. Multi-temporal comparability was ensured through consistency masking and leaf-off baseline initialization of empty space. Voxel transmittance and attenuation dynamics were analyzed across three areas of interest, covering a mixed plot, a beech and a Douglas fir patch. The results revealed distinct phenological signatures: European beech showed high-amplitude seasonal variation (70% attenuation volume reduction from summer to winter) with rapid spring foliation and stable winter baselines, whereas Douglas fir exhibited 12.7% variance with delayed growth onset and no stable winter baseline. Inter-annual comparisons indicated increasing blocking biomass with upward vertical shifts in transmittance profiles from tree growth and crown expansion. Two-dimensional attenuation mapping enabled the detection of individual tree dynamics and discrete structural disturbances. We demonstrate that consistent high-frequency UAV LiDAR acquisitions enable forest morphology monitoring through relative voxel transmittance. Our occlusion-aware framework differentiates structural changes from sampling biases, providing insights into forest dynamics at high spatio-temporal resolution.

**Graphical Abstract:** 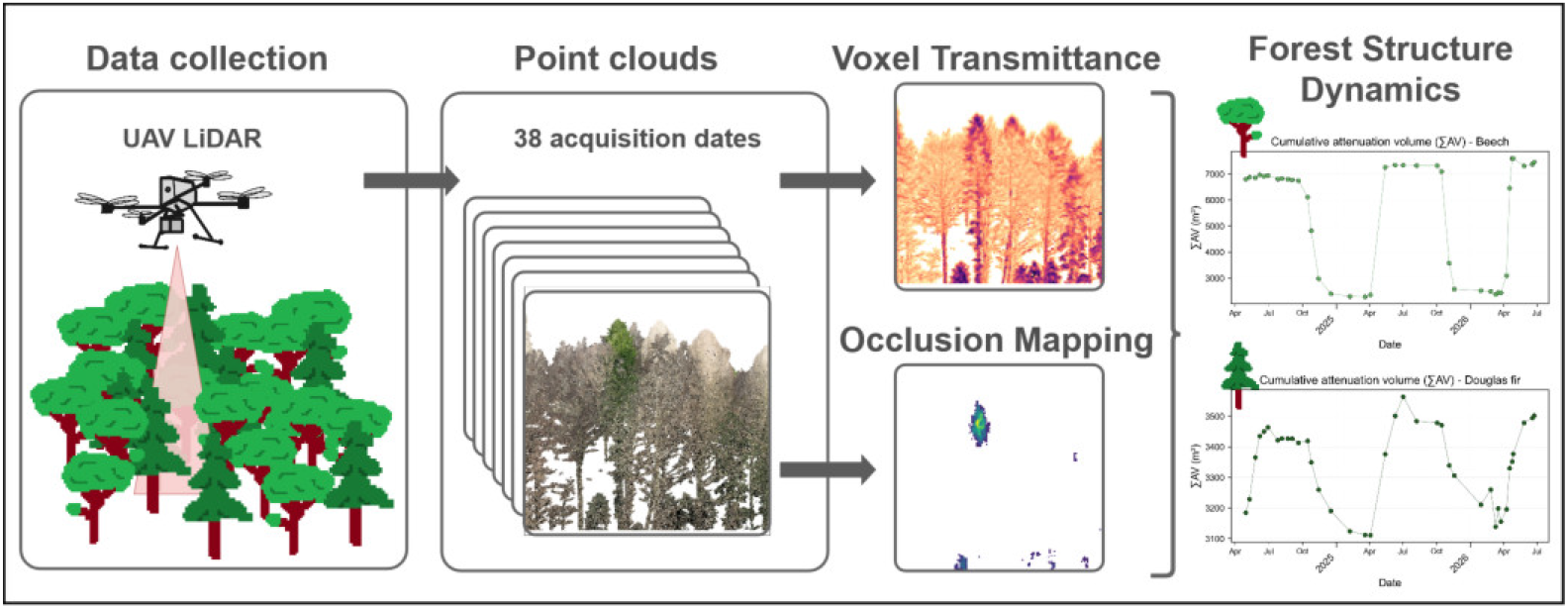

**Highlights:**

- High-frequency UAV LiDAR (38 flights) tracks forest structural dynamics *>* 2 years.
- Consistent acquisitions and occlusion-aware framework allow inter-comparability.
- Relative voxel transmittance is a robust proxy for monitoring 3D forest morphology.
- Beech showed 70% attenuation change, Douglas fir only 12.7% with delayed growth.
- 2D attenuation maps detect individual tree dynamics and structural disturbances.

## 1. Introduction

Forest structure varies in density, spatial arrangement and complexity and is a fundamental driver for ecosystem variables such as biomass, carbon sequestration and the microclimate. It influences light interception (Niinemets, 2010), wind velocity, gas exchange (Bienert et al., 2010), and the redistribution of precipitation through retention and throughfall (Pérez-Suárez et al., 2014), thereby impacting soil moisture (Tyagi et al., 2013), photosynthetic capacity, evapotranspiration, and the overall energy balance (Ellsworth and Reich, 1993; Norman and Campbell, 1989). Accordingly, these ecosystem variables are impacted by changes in forest morphology driven by phenology, growth, forest management and various disturbances (e.g. Frolking et al., 2009; Keenan et al., 2014). Common proxies for quantifying forest density include transmittance describing the fraction of incident light passing through the forest volume, or vice versa, the attenuation of light. From these, plant traits such as plant area density (PAD) or leaf area density (LAD) (Monteith and Unsworth, 1990) are commonly derived. However, mapping forest density and its magnitude of change in high-resolution and 3D remains a non-trivial task.

Manual measurement methods are expensive and time-consuming and knowledge on spatio-temporal structure dynamics remains limited with field methods such as destructive sampling, empirical allometry functions (Albrektson, 1984; Grier and Waring, 1974), litter-traps (Cutini et al., 1998), and the inclined point-quadrat method (Wilson, 1959, 1960). Therefore time-of-flight (ToF) light detection and ranging (LiDAR) has become a standard for capturing forest structure (Koch, 2010). While studies demonstrated potential for full-waveform LiDAR to retrieve vegetation density via radiative transfer models (RTMs), LiDAR data is commonly processed into point clouds of discrete-return point representing the explicit 3D structure.

Research often focuses on turning these point clouds into metrics to estimate transmittance and quantify forest density. This is achieved via linking beam passing probability to the point-quadrat method (e.g. Béland et al., 2011; Morsdorf et al., 2006) or to Beer-Lambert’s law (Monsi, 2004; Nilson, 1971; Ross, 1981). These methods are based on gap fraction metrics defining the probability of a light ray passing through the canopy without intersection, a value that can be estimated via the emission of millions of LiDAR pulses.

Gap fraction metrics have evolved from simple return ratios (Almeida et al., 2019; Bouvier et al., 2015; Morsdorf et al., 2006) to more complex models that account for pulse power attenuation of fractional returns (Arnqvist et al., 2020; Fleck et al., 2012; Solberg et al., 2009). Recent advancements incorporate the optical path length through turbid media via ray-tracing (Grau et al., 2017) or path length distributions (Pimont et al., 2018, 2019; Soma et al., 2021) to correct for scanner properties and viewing angles. A holistic, path-length-independent method was proposed by Vincent et al. (2017), which integrates weighted pulse fractions and the diverging LiDAR footprint, normalized by the mean traversed path length.

Driven by computational advancements and increased availability of high-spatially resolved sensor data, forest density mapping has shifted from 2D aggregated distributions (e.g. Almeida et al., 2019; Bouvier et al., 2015; Morsdorf et al., 2006; Véga et al., 2016) to 3D spatially explicit voxel mapping (e.g. Arnqvist et al., 2020; Grau et al., 2017; Pimont et al., 2018; Schneider et al., 2014; Vincent et al., 2017), enabling the investigation of biomass and phenology at unprecedented resolutions.

Yet, studies investigating density dynamics are scarce, partly because consistent multi-temporal LiDAR datasets are rare. Many studies are limited to bi-temporal comparisons (e.g. Bienert et al., 2018; Lin et al., 2022; Nguyen et al., 2020; Niwa, 2025). While some have expanded to 3–7 acquisitions (Gretler et al., 2026; Matasci et al., 2018; Slavík et al., 2020; Winstanley et al., 2024; Xue et al., 2026; Yin et al., 2024), high-resolution temporal monitoring is usually limited to static installations (Calders et al., 2023; Campos et al., 2021; Korpelainen et al., 2026; Shcherbacheva et al., 2024). Extensive high-frequency collections remain rare; for instance, Calders et al. (2015) utilized 48 terrestrial laser scanner (TLS) scans over 20 weeks to monitor spring phenology, while recent UAV laser scanning (ULS) datasets from Botticelli et al. (2026) spanning the autumn season of two years have enabled the analysis of silvicultural impacts on autumn phenology (Bajocco et al., 2026).

However, the transition to temporal monitoring reveals a critical dependency on sensor properties and acquisition settings, where variations in wavelength, sensor sensitivity, and beam divergence can introduce systematic biases (Gretler et al., 2026; Vincent et al., 2023). Vincent et al. (2023) showed up to 25 % differences in attenuation profiles of different sensors, while Bartholomeus et al. (2026) and Gretler et al. (2026) highlight the influence of flight geometry and sampling design. Varying viewpoints, scan angles, flight height and sampling densities alter optical path lengths, the amount of observed and occluded areas, footprint size and at-canopy pulse power (Almeida et al., 2019; Brede et al., 2022; Gassilloud et al., 2025; Shao et al., 2019) further complicating the retrieval of consistent density metrics. Because gap fraction metrics are particularly prone to these inconsistencies, it becomes nearly impossible to distinguish true structural dynamics from methodological artifacts. Consequently, while statistical inter-calibration is possible (Shao et al., 2019; Vincent et al., 2023), the gold standard for reliable multi-temporal analysis remains the use of identical sensors, harmonized settings and path-length-corrected methods (see Gretler et al., 2026; Vincent et al., 2017).

A persistent bottleneck remains the validation of derived PAD and LAD values. Destructive sampling is rarely feasible at scale, leaving studies to rely on virtual simulations (Pimont et al., 2018) or field methods for local calibration like digital hemispherical photography (DHP), litter-traps and LAI-2200 (e.g. Morsdorf et al., 2006; Solberg et al., 2009; Vincent et al., 2017) which lack 3D resolution and introduce their own biases. Ambiguity is added by assumptions of homogeneous vegetation requiring clumping correction and leaf angle distributions which depend on tree species, environmental factors and undergo sub-daily changes (e.g. Kremer et al., 2026). In contrast, while LiDAR-derived transmittance may not represent an absolute physical value, it is a direct measurement of the system’s ability to penetrate the canopy. By maintaining consistent acquisition parameters, changes in observed transmittance can serve as a more robust proxy for shifts in forest morphology than absolute PAD estimations, effectively bypassing the layers of ambiguity. This study aims to quantify high-frequency, consistent, and occlusionaware forest structural density at the plot scale within a European mixed temperate forest. Therefore an extensive multi-temporal ULS acquisition of 38 flights was conducted over slightly more than two years, starting in April 2024 and ending in June 2026. Voxel transmittance is calculated following Vincent et al. (2017, 2021) using a custom Python implementation for handling large datasets. This work demonstrates the potential of using relative changes in voxel transmittance as a direct proxy for dynamic shifts in forest morphology. Specifically, we investigate the temporal evolution of voxel transmittance and occlusion through four primary objectives: (1) quantifying temporal structural dynamics at the plot level, (2) analyzing variations in vertical height profiles, and (3) mapping 2D spatial heterogeneity to identify hotspots of structural change. Furthermore, we assess species-specific structural dynamics by comparing patches dominated by European beech and by Douglas fir.

## 2. Material and methods

### 2.1. Study site

This study was conducted at the recently established ECOSENSE field site, which is a low-impact managed, mixed temperate forest located in the southwest of Germany in the upper Rhine valley that was equipped with dense sensor-network for monitoring forest processes (Werner et al., 2024). It is located at an altitude between 491 m to 525 m (EPSG:32632) on the ridge of a hill with a terrain slope between 0^◦^ and 30^◦^. Focus was set on the central section of the field site, which covers about 8 ha (see Fig. 1).

**Figure 1.**
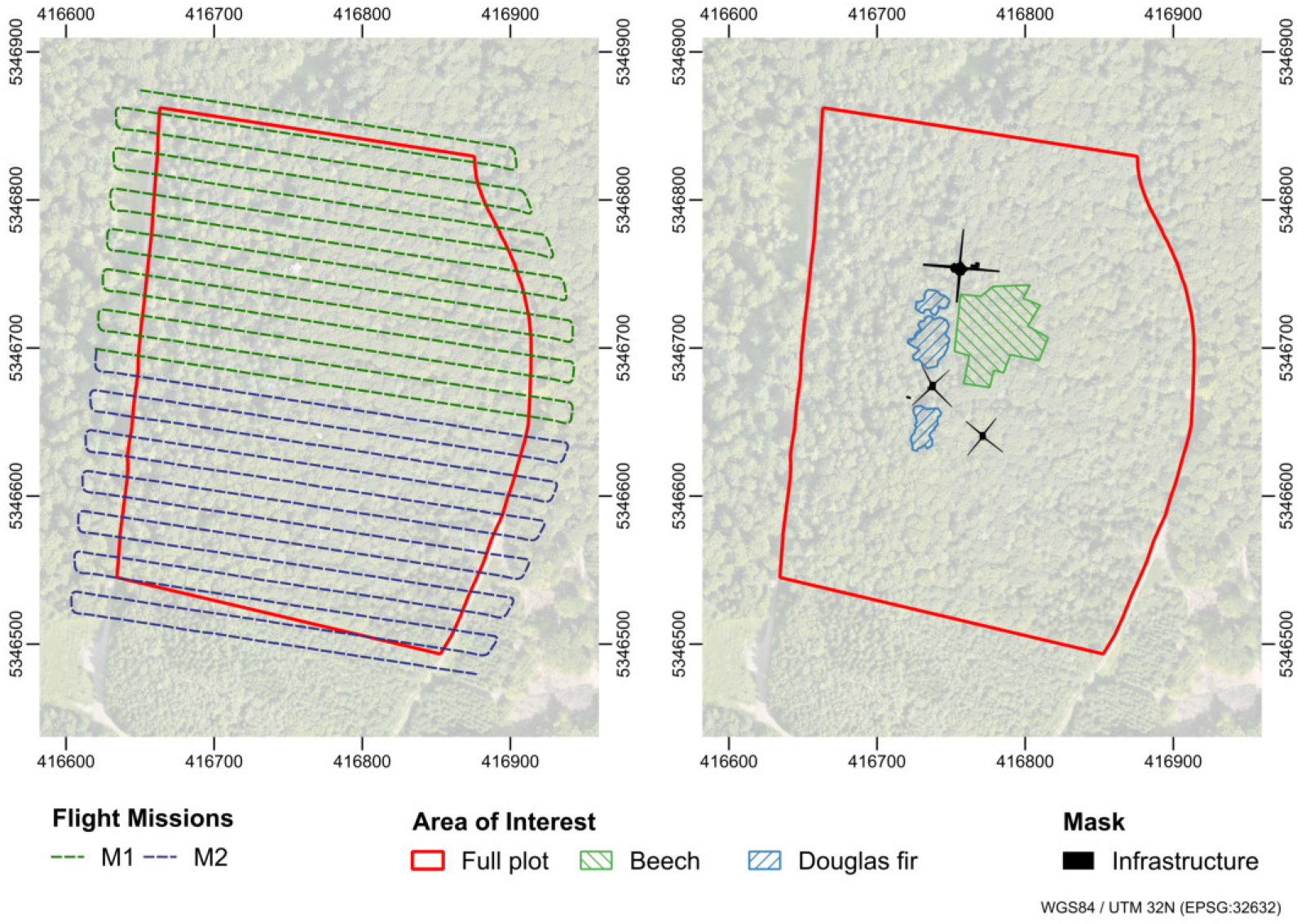
Study site with trajectory of the two flight missions (M1 and M2). Area of interest (AOI) covering the full plot, beech and Douglas fir patches. Human-built structures are masked for analysis.

The full plot is dominated by European beech trees (*Fagus sylvatica*) intermixed with patches of Douglas fir (*Pseudotsuga menziesii*), European silver fir (*Abies alba*) and dispersed trees of English oak (*Quercus robur*), Scots pine (*Pinus sylvestris*) and European larch (*Larix decidua*) (Werner et al., 2024). The forest structure is mostly single layered with only little midand understory vegetation. Tree height reaches up to about 41 m in the full plot (March 2023). Human-built structures located in the full plot include three canopy access towers and two containers for equipment storage (a detailed characterization of the field site can be found in Tesch et al., 2025). Three area of interest (AOI) are defined for this analysis: one covering the full plot, and two covering mostly homogeneous beech and Douglas fir patches to investigate species-specific dynamics (see Fig. 1). Two exemplary pictures of the beech and Douglas fir patch are shown in Fig. A.13.

### 2.2. Data Acquisition

Data acquisition was conducted using a DJI Matrice 300 RTK UAV and a DJI Zenmuse L2 LiDAR scanner. The DJI Zenmuse L2 LiDAR scanner operates at a wavelength of 905 nm and has an integrated inertial measurement unit (IMU). The laser beam divergence is specified with 0.2 mrad horizontal * 0.6 ^◦^ vertical, resulting in a footprint size of 2.4 cm * 7.2 cm at a sampling distance 60 m. The LiDAR scanner can capture up to 5 returns with a sampling rate of 240 kHz (for more details on the specifications see DJI, 2024).

Flight trajectories were created with UgCS (SPH Engineering), and scanner settings were manually adapted on the controller. The flight missions were planned considering flight duration, the number of batteries, the desired area coverage, and optimized based on the insights gained by Gassilloud et al. (2025). In this study, we used the Lissajous scanning mode, a so-called nonrepetitive pattern with a field of view (FOV) of about 70 ^◦^ (horizontal) * 75 ^◦^ (vertical). The sensor tilt angle was set to face nadir. The field site geometry and topography favored a west-east flight pattern to maintain connection to the remote controller.

The sampling design aimed to increase the number of viewpoints, sampling density and thus reduce occlusion (Gassilloud et al., 2025; Kükenbrink et al., 2025). The horizontal distance between flight paths corresponded to 50% overlap at the top of canopy (∼ 40.5 m height) with the sensors FOV, as recommended in (Kükenbrink et al., 2017). The flights were performed at comparably low flight heights of about 60 m above ground following a digital terrain model (DTM) with flight speeds around 5 ms^−1^ (see Table 1). The flight mission was split into two parts (M1 and M2) to allow for changes of batteries in between.

**Table 1.** Flight mission parameters for M1 and M2. To ensure comparability between acquisitions, they were kept constant for all flights.

| Mission | Flight<br>height (m) | Flight speed<br>($\text{ms}^{-1}$) | Azimuthal flight<br>direction (degree) | Horizontal trajec-<br>tory distance (m) |
| --- | --- | --- | --- | --- |
| 1, 2 | 60 | 5 | 98 | 13.68 |

For precise geo-referencing and flight path routing a DJI D-RTK 2 GNSS mobile reference station was placed on a pre-determined fixpoint. IMU calibration was performed before and after each flight. All flights were performed under nearly windless weather conditions to minimize branch movement and improve inter-flight comparability. Data collection was conducted with variable time intervals starting from 30 April 2024 to 23 June 2026, resulting in a time series of 38 flight days. Intervals between flight days were heavily determined by equipment capacities and weather conditions, whereby autumn and winter seasons were often restricted by fog. The raw data of all flights was processed using DJI Terra software (v. 4.3) to generate point clouds projected to WGS84/ UTM 32N (EPSG:32632). As reconstruction algorithms turned out to vary significantly and produce different point clouds between individual versions (see DJI, 2026), we decided to commit to one common version for all flights to achieve best inter-flight comparability.

Not all conducted flights were successful. Sometimes RTK positioning or IMU errors became apparent only during processing, and point clouds could not be (fully) constructed. In these cases, PPK was used to successfully reconstruct the point clouds. For PPK processing, a (virtual) reference station provided GPS correction data and was used to replace the D-RTK base station. An overview of the flights including respective reference stations is provided in Table B.2. Resulting point clouds consist of discrete returns labeled with the return number, the number of returns, and the GPS time of the LiDAR beam emission.

### 2.3. Pre-processing

To track LiDAR beams through a voxel grid with a ray-tracing algorithm, it was necessary to reconstruct their trajectory from the position of pulse emission to the last return. Following Gassilloud et al. (2025), the LiDAR sensor positions trajectories with GPS time stamps were reconstructed (50 positions per second) from the point clouds.

Even though most datasets were generally well aligned (within the tolerance of error propagation by base station set-up, D-RTK, LiDAR and IMU precision or PPK processing), small spatial offsets were observed within the time series. Therefore, the flight mission M1 acquired on 7th February 2025 was used as a reference dataset, and all datasets were manually aligned in cloud compare (v. 2.13.2). Both flight missions covered a canopy access tower, on which the point clouds were visually matched, resulting in spatial shifts in the range of ∼ 0-12 cm on the three axes. The transformation matrix was stored and applied to the point cloud and the reconstructed sensor positions before further processing.

The point clouds of both flight missions (M1, M2) from the same day were merged into a single file, and the sensor position trajectories were concatenated. For each individual LiDAR beam, the pulse origin was estimated with linear interpolation using their GPS time and the chronologically previous and following reconstructed sensor position (Gassilloud et al., 2025). For simplicity, this was done for single and multiple return LiDAR beams. Point clouds were filtered to ensure consistent and robust results. Discrete returns of individual beams were determined and the relevant scalar fields examined: We identified and removed LiDAR beams and corresponding returns with inconsistent “return number” labeling, inconsistent “number of returns” labeling and an inconsistent distance between pulse origin and corresponding returns (needs to be strictly increasing). As a result, around ∼ 0.3-0.7 % of the LiDAR beams and their corresponding returns were removed.

### 2.4. Transmittance calculation

The voxel transmittance was calculated following the methodology described in Vincent et al. (2017). A custom python implementation was developed, optimized for handling larger datasets compared to the current Amapvox implementation (v. 2.2.1). The code is accessible via the following repository: https://github.com/MGEOS/CANOPy.

The initial formula by Vincent et al. (2017) was adapted by Vincent et al. (2021), where the normalized voxel transmittance *P_Gap_* is calculated by solving Eq. 1:

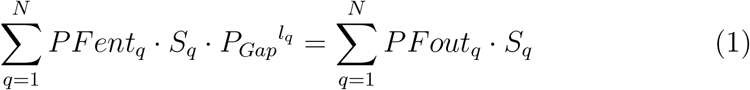

where:

*P_Gap_* = transmittance per unit length optical pathway

*PFent_q_* = entering pulse fraction of pulse *q* (energy)

*PFout_q_* = exiting pulse fraction of pulse *q* (energy)

*S_q_* = beam section of beam *q* at voxel center

*l_q_* = theoretical beam length of beam *q* in voxel

*N* = number of beams entering the voxel

The ray tracing applied in Gassilloud et al. (2025) was adapted to map traversing beams and respective properties to each voxel. While Vincent et al. (2017) assigned averaged return intensity fractions based on the return number and number of returns, this study used equal weighted fractions, as limited information was available regarding the processed intensity values provided by the DJI terra software. For a beam *q* with *n_all_* returns, each remaining return was considered to contribute 1*/n_all_* to *PFent_q_* and hence *PFout_q_* = *PFent_q_* − (*n_intercepted_*∗ 1*/n_all_*). The non-linear equation (Eq. 2) has no analytical solution, and *P_Gap_* had to be estimated. Therefore, an objective function *f* (*P_Gap_*) was defined, which represents the error of any estimated *P_Gap_*. We searched for *P_Gap_* so that *f* (*P_Gap_*) = 0.

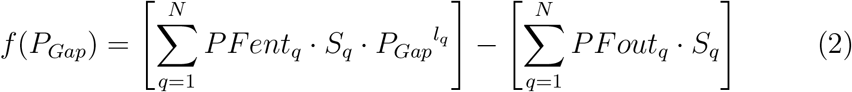

With *N* being the total number of LiDAR beams passing the voxel and therefore *PFent_vox_* = {*PFent*_1_*, …, PFent_N_* } being an array of the entering pulse fractions, *PFout_vox_* = {*PFout*_1_*, …, PFout_N_* } the corresponding exiting pulse fractions, *S_vox_*= {*S*_1_*, …, S_N_* } the corresponding beam sections at voxel center and *l_vox_* = {*l*_1_*, …, l_N_* } the corresponding theoretical path lengths within the voxel.

To estimate *P_Gap_*for a voxel, we used a bisection algorithm (see Alg. 1). We set a lower bound *a* = 0 and an upper bound *b* = 1 and initialize *P_Gap_* = (*a* + *b*)*/*2. Now we could evaluate *f* (*P_Gap_*). If *f* (*P_Gap_*) was close to 0 (here *<* 10^−8^), we found the root. Else, we decreased the search interval by re-defining the boundaries. If *f* (*P_Gap_*) *<* 0, the root was to the right and we set *a* = *P_Gap_*. If *f* (*P_Gap_*) *>* 0, the root was to the left and we set *b* = *P_Gap_*. In the next iteration, we retrieved a new estimate *P_Gap_* = (*a* + *b*)*/*2 with the new boundaries, converging *P_Gap_* towards the true value. Similar to the implementation in Amapvox, when 200 iterations were reached without *f* (*P_Gap_*) *<* 10^−8^, we took *P_Gap_* = (*a* + *b*)*/*2 as our final estimate.

The voxel size has a strong impact on absolute transmittance values and occlusion mapping. It ideally balances the available sampling density and LiDAR beam footprint size while enabling highly spatially resolved mapping for a meaningful analysis (Béland et al., 2011; Kükenbrink et al., 2017, 2025). Here, we used a voxel size of 0.25 m to account for GPS accuracy, coregistration, branch bending, and tree movement, while maintaining a high spatial resolution, which aligns with previous studies for high sampling density datasets (Grau et al., 2017; Schneider et al., 2019). The voxel space was normalized with a digital terrain and reduced to a slice ranging from 1 m to 41 m height above ground for further analysis.

#### Algorithm 1 Pseudocode of transmittance estimation after Vincent et al. (2021) with bisection algorithm.

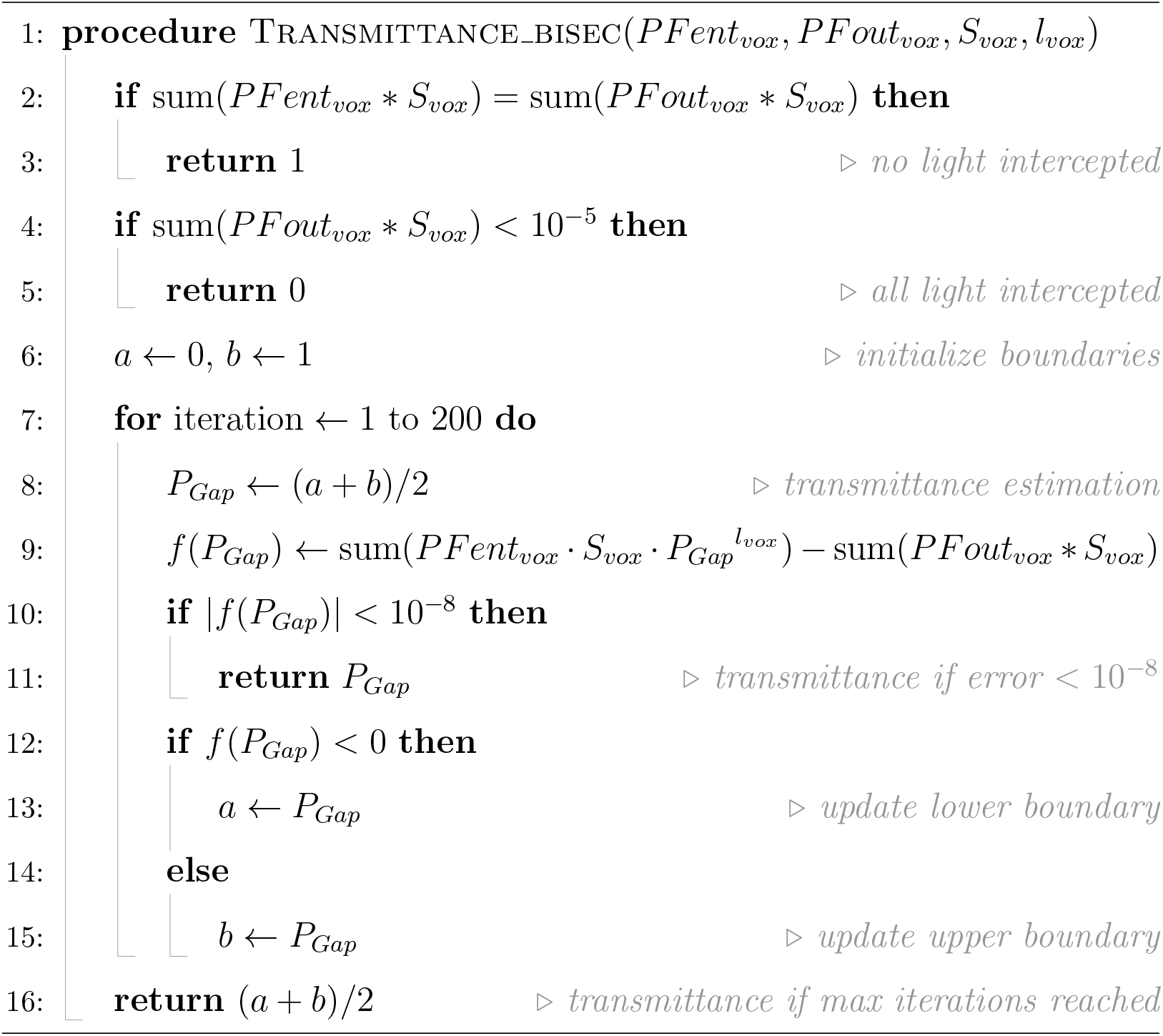

### 2.5. Post-processing

To account for occlusion effects, we additionally classified the voxel space into four categories: empty, occluded, filled and unobserved, following the same approach as in Gassilloud et al. (2025).

Due to the sampling density achieved in this study, unobserved voxels were nonexistent within the canopy. However, occlusion remained a significant challenge, as it varied strongly within the phenological cycle (Gassilloud et al., 2025; Kükenbrink et al., 2017). In summer, under leaf-on conditions, large proportions of ULS LiDAR beams may be blocked, resulting in occlusion within and below the canopy. In contrast, winter (leaf-off) acquisitions have less occlusion and allow for a better quantification of the actual observable volume.

In the analysis of multi-temporal LiDAR data which covers different stages of phenology, there is a high risk of monitoring change that results from sampling different areas due occlusion effects and not due to actual structural change. This problem propagates to all attempts of filling occluded space with interpolation methods. We therefore aggregate a mask from all datasets to define which voxels to include and to exclude for further analysis and thus ensure consistency across the time series. Since the likelihood of a voxel being occluded at least once in 38 datasets is high, we had to choose a criterion that did not exclude too many voxels while maintaining comparability between the datasets (see Fig. C.14).

Therefore, we applied following rules: (1) Voxels that were always occluded throughout the time series were excluded from the analysis. (2) All voxels occluded *>* 3 times throughout the time series were also excluded from the analysis. (3) If a voxel was occluded ≤ 3 times and was never unobserved through the time series, they might be considered for analysis. In this case, they were excluded from analysis when occluded, otherwise they were considered. We re-classified all excluded voxels as occluded voxels and discarded them for the analysis of transmittance values.

Kükenbrink et al. (2017) showed that most occluded space in airborne datasets acquired with leaf-on condition is actually empty space. To counteract occlusion even further, we divided the volume into space that may contain plant material throughout the time series and space that was likely empty.

We took the reference leaf-off winter acquisition from 7th February 2025 and created a spatial buffer of 1 m around all voxels that were not empty (hence are filled, occluded or unobserved and thus may contain plant material). The remaining volume could be regarded as “true empty space” and transmittance was set to 1 across the time series, even though these voxels might have had been previously excluded. Fig. C.15 shows the mean occluded volume across all dates after the application of exclusion rules and initialization of empty space, highlighting where areas excluded from analysis and thus where forest density is likely underestimated. Finally, human-built structures (see Fig. 1) such as the canopy towers were masked manually and excluded from the analysis.

### 2.6. Analysis

With data processing, we obtained for each date a voxel space with transmittance values (*P_Gap_*) that provided a direct measurement of canopy penetrability, including a classification and mapping of occluded voxels. To provide a more intuitive representation of light blocking biomass, we also calculated the voxel attenuation (*A* = 1 − *P_Gap_*) as the inversion of transmittance. As a proxy for the distribution of light-blocking biomass, we derived the attenuation volume (Σ*AV* = *A* ∗ *V_V ox_*) by multiplying the attenuation of each voxel by its voxel volume (*V_vox_* = 0.25 *m*^3^). The absolute values of transmittance and (cumulative) attenuation volume are primarily used for relative comparison across the time series. The occluded volume (*V_occ_*) was derived as the fraction of occluded voxel volume relative to the total voxel volume below the top of canopy. These metrics were aggregated across different scales to address the studies primary objectives.

On plot level, we tracked the temporal trajectories of the mean attenuation of filled voxels (*A_filled_*) in the three AOIs to monitor vegetation density, the cumulative attenuation volume (Σ*AV*) as a proxy for total blocking biomass, and the occluded volume fraction to highlight the impact of phenology on data consistency.

To analyze vertical height profiles for the three AOIs, we aggregated the mean transmittance (*P_Gap_*) and the cumulative attenuation volume for each date along vertical voxel layers.

To map the 2D distribution of structural change, we created cumulative attenuation maps for each date with the attenuation volume (Σ*AV*) summed along the vertical voxel axis. Further, we investigated an exemplary bitemporal difference between a summer (23 June 2026) and winter (25 February 2025) data acquisition, and the relative amplitude (*RAV* = (*AV_max_*− *AV_min_*)*/AV_mean_*) as the normalized structural change of the time series.

## 3. Results

The repeated ULS acquisitions resulted in an extensive multi-temporal collection of point clouds covering more than two years. For each date, a 3D voxel space was derived with estimated transmittance values and with an occlusion classification. Fig. 2 shows an exemplary subset of the three resulting datasets. All data used in this study is made available [HERE - link will be updated].

**Figure 2.**
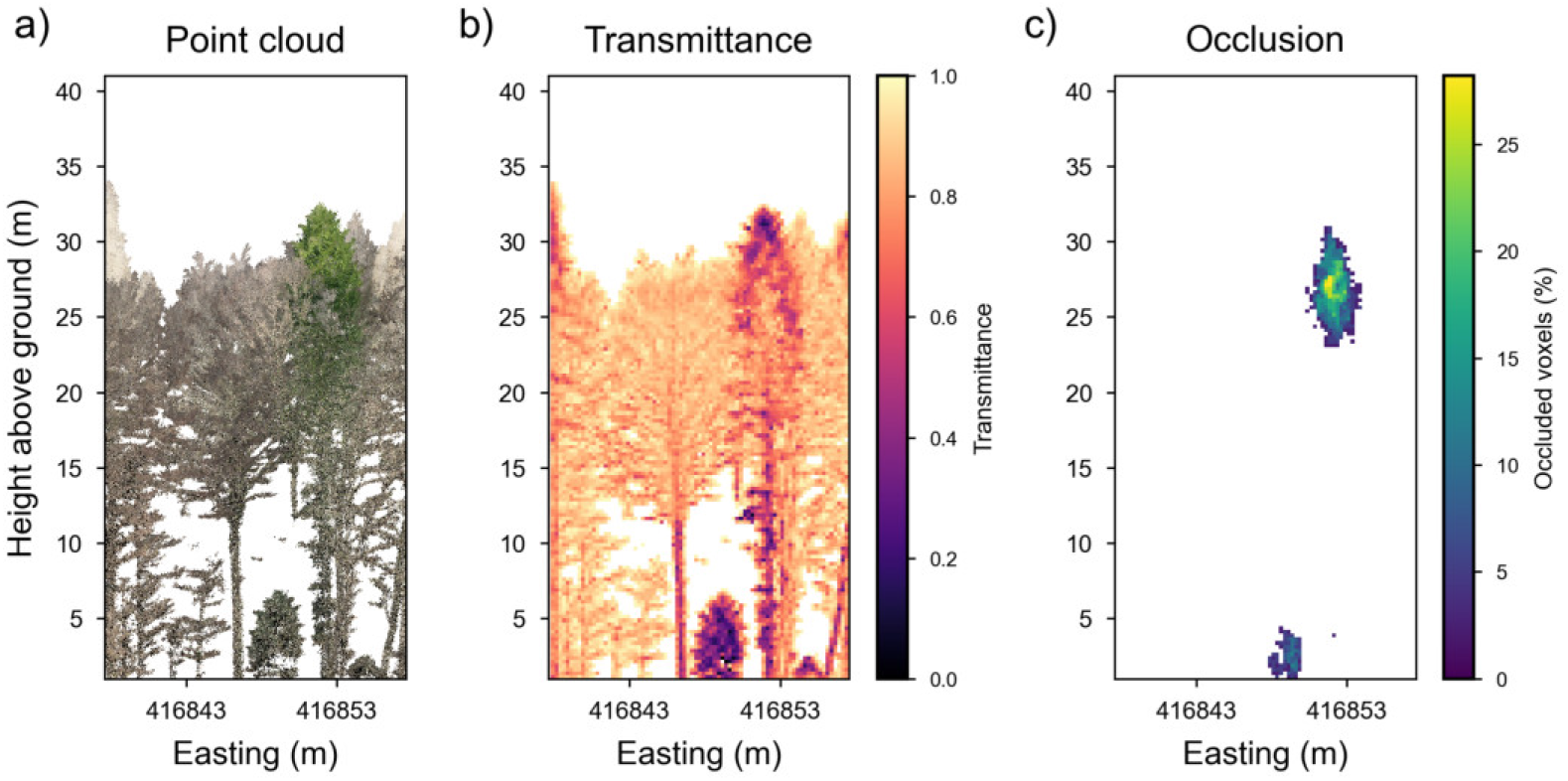
A representative subset from 7 February 2025 covering a transect of 10 m (north) ∗ 20 m (east), aggregated along the northing axis. It shows a) the point cloud, b) the mean transmittance of filled voxels, and c) the percentage of occluded volume.

The following results present the quantification of temporal variations in transmittance, attenuation as well as occluded voxel volume, structured by spatial aggregation levels to address our primary objectives: First, we analyze the overall dynamics at the individual AOI level (subsection 3.1), followed by an examination of vertical height profiles (subsection 3.2), and finally, the generation of 2D maps to characterize the spatial heterogeneity of the full plot (subsection 3.3).

### 3.1. Temporal dynamics on plot level

To analyze changes in vegetation density, we investigated the temporal dynamics at AOI level by tracking the mean attenuation of filled voxels, the cumulative attenuation volume and occluded volume below top of canopy over time.

#### 3.1.1. Mean attenuation of filled voxels over time

Fig. 3 shows the mean voxel attenuation of filled voxels over time for the three AOIs. Fig. 3a) shows the mean voxel attenuation of filled voxels (*A_filled_*) in the beech patch, which underlies strong seasonal fluctuations. The time series starts on 30 April 2024 close to the end of foliation. *A_filled_* peaked with 0.5 on 26 May 2024, before entering a phase of slow degradation during summer. This gradual decline is likely attributed to the loss of biomass due to environmental factors while lacking the formation of new leaves. A rapid decline is observed on 15 October 2024, corresponding to autumn senescence, with a stable leaf-off baseline around ∼ 0.14 reached on 17 December 2024.

**Figure 3.**
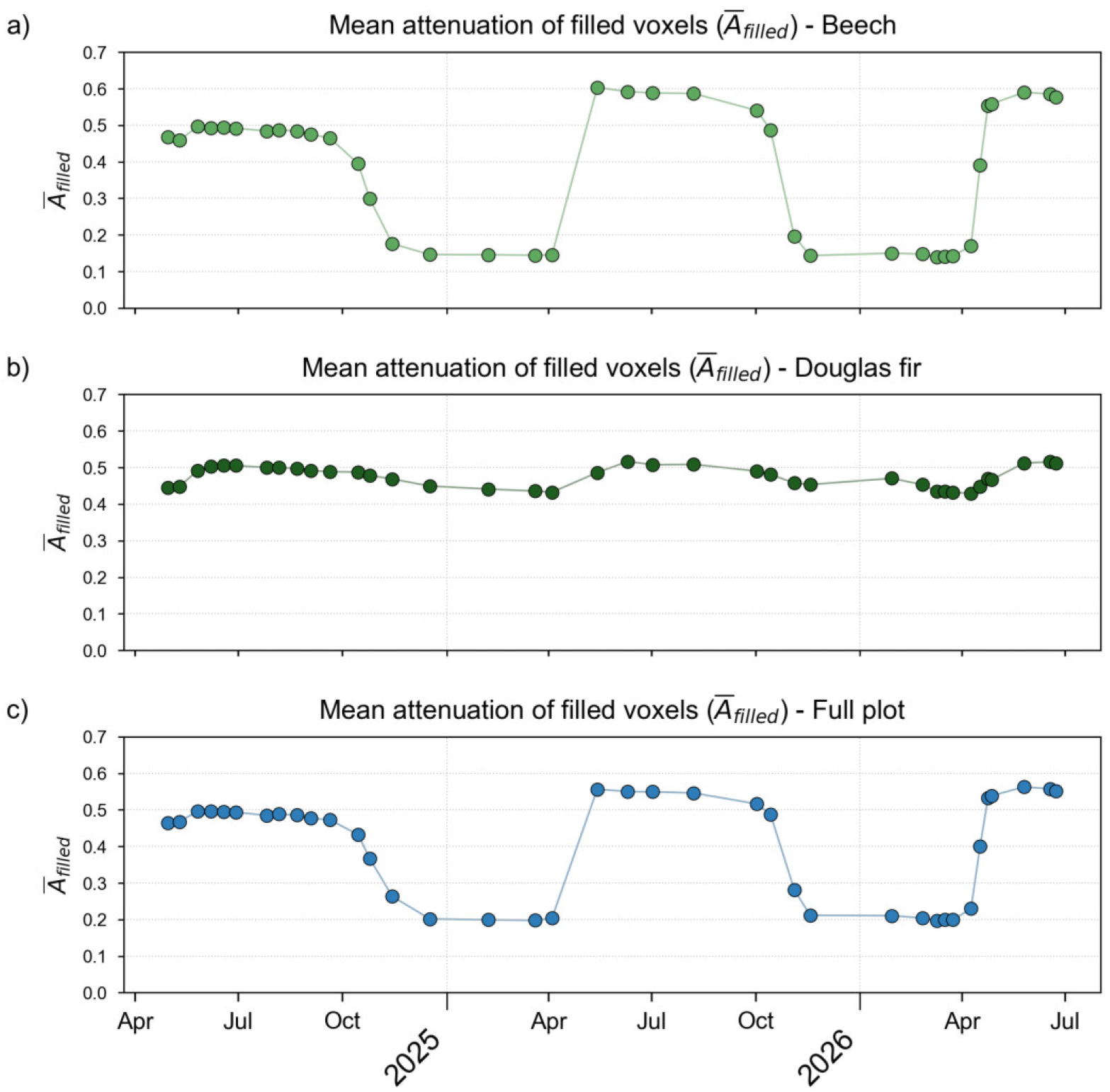
Mean attenuation of filled voxels in the three AOIs: a) beech patch with strong seasonal variations, b) Douglas fir patch with less pronounced phenology, and c) full plot, dominated by beech dynamics.

In 2025 bud burst occurred between 4 April and 14 May, followed by a higher peak attenuation (0.6) than in the previous summer, suggesting an overall increase in forest density. The same phenological pattern continues with a similar winter baseline (0.14) and a very rapid increase in attenuation during foliation between 9 April and 24 April 2026. The next maximum (0.59) is reached on 26 May 2026, albeit without an increase towards 2025.

Also the Douglas fir plot in Fig. 3b) shows seasonal patterns, even though with significantly lower variance. In 2024, the strongest increase in *A_filled_*starts after 10 May 2024, which is later than the beech patch where foliation is almost completed. The summer peak attenuation (0.51) is reached on 18 June 2025. After maxima, attenuation degrades until the following growth period, with an increased decline during autumn. In contrast to beech, no stable winter baseline is observed. Similar behavior is observed at the start of 2025, where peak attenuation with ∼ 0.52 is slightly higher.

A sudden increase in *A_filled_* is observed on 29 January 2026, followed by a measurement on 25 February 2026 at a level similar to that of 18 November 2025. Again degradation continues until the start of new growth period. The following summer *A_filled_*maximum in 2026 is similar to 2025.

The dynamics of the full plot in Fig. 3c) are heavily dominated by beech trees. Here, we can also see a stronger increase of maximum *A_filled_* in 2024(0.5) to summer 2025 (0.56), and a similar maximum in 2026 (0.56).

#### 3.1.2. Cumulative attenuation volume over time

While the mean attenuation describes the intensity of blocking within filled voxels, the cumulative attenuation volume (Σ*AV*) shown in Fig. 4 accounts for the total extent of the blocking biomass across the AOIs.

**Figure 4.**
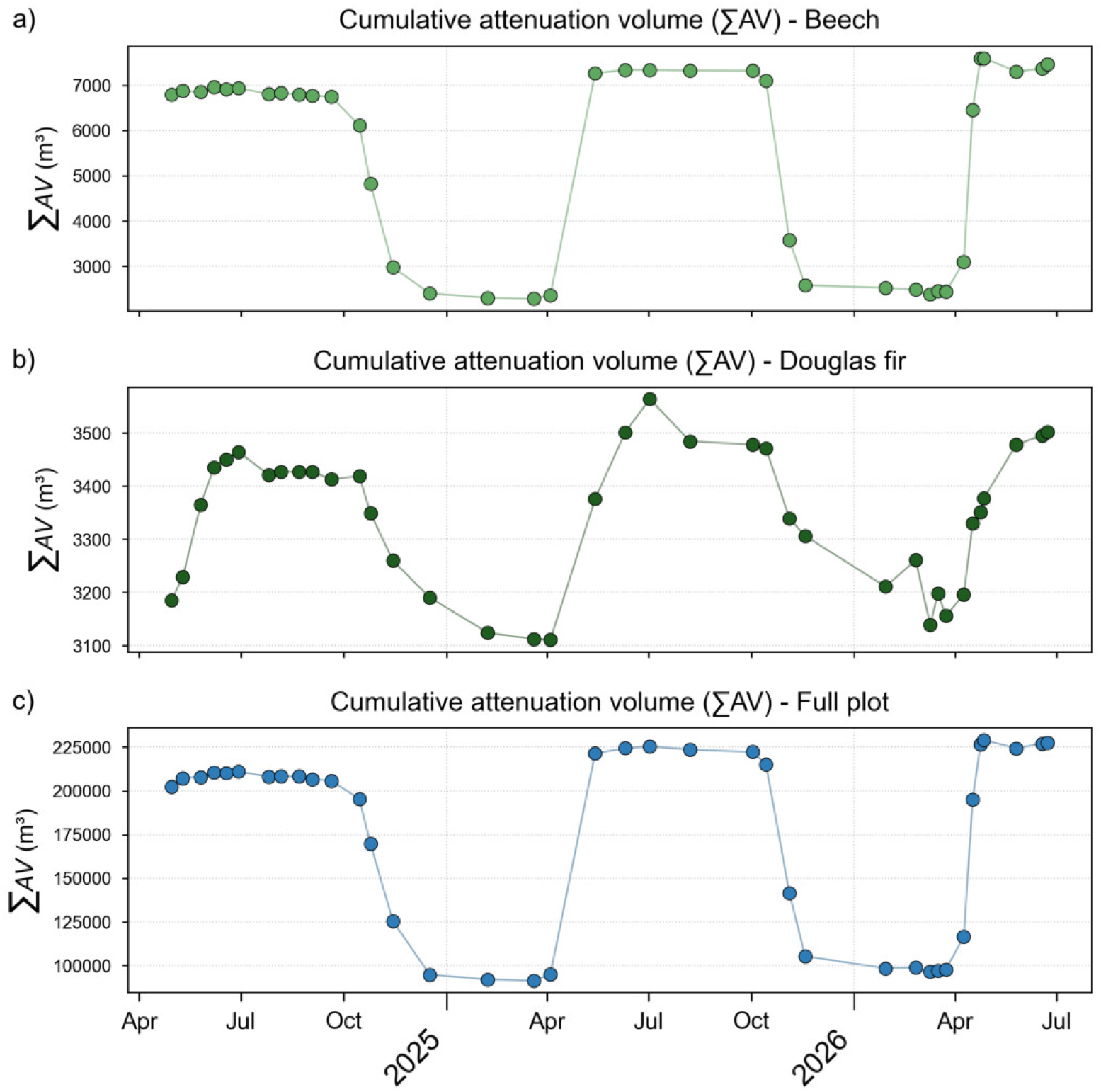
Cumulative attenuation volume of the three AOIs: a) beech patch with 70 % reduction relative to the maximum, b) Douglas fir patch with 12.7 % reduction, and c) full plot with 60,2 % reduction.

The beech patch in Fig. 4a) shows strong seasonal variations in blocking biomass that closely aligns with the trends observed for mean attenuation of filled voxels (see Fig. 3). In 2025, the growth period starting in April led to a higher summer plateau of blocking biomass compared to 2024, indicating an overall increase of blocking biomass. Compared to *A̅_filled_*, Σ*AV* shows a slight increase during summer 2026 as well. While *A̅_filled_* and Σ*AV* follow a similar decline during summer 2024, Σ*AV* remains on a more constant summer plateau in 2025, while *A̅_filled_* slowly decreases. The difference between the maximum and minimum Σ*AV* corresponds to a 70 % reduction relative to the maximum.

Also the Douglas fir patch in Fig. 4b) follows closely the trend of *A̅_filled_*. However, after maximum Σ*AV* is reached, attenuation suddenly drops, followed by a stable summer plateau, both in 2024 and 2025. Σ*AV* decreases rapidly during autumn, stabilizing shortly before growing season starts again. Also here maximum Σ*AV* in 2025 increased towards 2024, however the peak in 2026 is likely not captured by the end of this dataset. The difference between the maximum and minimum Σ*AV* corresponds to a 12.7 % reduction relative to the maximum. Also here the cumulative attenuation volume exhibits strong variations in early 2026, including a sudden increase on 25 February.

The maxima in Σ*AV* for Douglas fir are again later than the maxima for beech. While the onset of growth is the same for beech and Douglas fir among the dataset, the increase of Douglas fir Σ*AV* spans a longer period of time. The start of sudden decline during autumn is again the same for beech and Douglas fir in the dataset. Yet, the decline of Douglas fir continues until next growing season while beech decreases faster and stabilizes during winter.

For the full plot in Fig. 4c), seasonal dynamics are again similar to the beech patch. The increase in summer maxima from 2024 to 2025 and the higher winter minimum in 2026 were more pronounced in the full plot than in the beech patch. The difference between the maximum and minimum Σ*AV* corresponds to a 60,2 % reduction relative to the maximum.

#### 3.1.3. Occluded volume over time

To highlight the impact of sampling bias, we analyzed the percentage of occluded volume (*V_occ_*) relative to the total volume below the top of the canopy for each AOI. Fig. 5 compares the inherent occlusion of the original datasets to the occlusion after applying the consistency mask and initialization of empty space (see subsection 2.5). In Fig. 5a), occlusion in the original datasets of the beech patch exhibits strong seasonal dynamics. While occlusion was negligible in summer 2024, it increased sharply in the subsequent years, exceeding 10.2 % in summer 2025 and reaching over 12.4 % in 2026. This indicates that canopy gaps closed as the canopy grew, which significantly limited the LiDAR’s penetration capability. After masking, variations in occlusion were substantially reduced. The occlusion remains relatively stable around ∼ 7.8 %, with smaller peaks during the summer of 2025 (9.4 %) and 2026 (10 %). The reduction towards the original datasets in summer is primarily attributed to the initialization of empty space, which reclassified a fraction of the occluded volume as empty. The Douglas fir patch in Fig. 5b) shows much less occlusion compared to the beech patch. Original occlusion values ranged from a minimum of 0.23 % to a maximum of 1.37 % in summer 2026. The structure of Douglas fir trees is dense; however, the patch is not densely populated and leaves gaps for below canopy exploration. Even though the absolute magnitude of change is small, the dynamics still follow the general phenological trend observed in mean attenuation and cumulative attenuation volume, with an increase in occlusion over the years. By the end of 2025 and beginning of 2026 a slight increase in occlusion is observed. Although starting already by end of 2025, the dynamics align with the dynamics observed for *A̅_filled_* and Σ*AV* (see Figs. 3 and 4).

**Figure 5.**
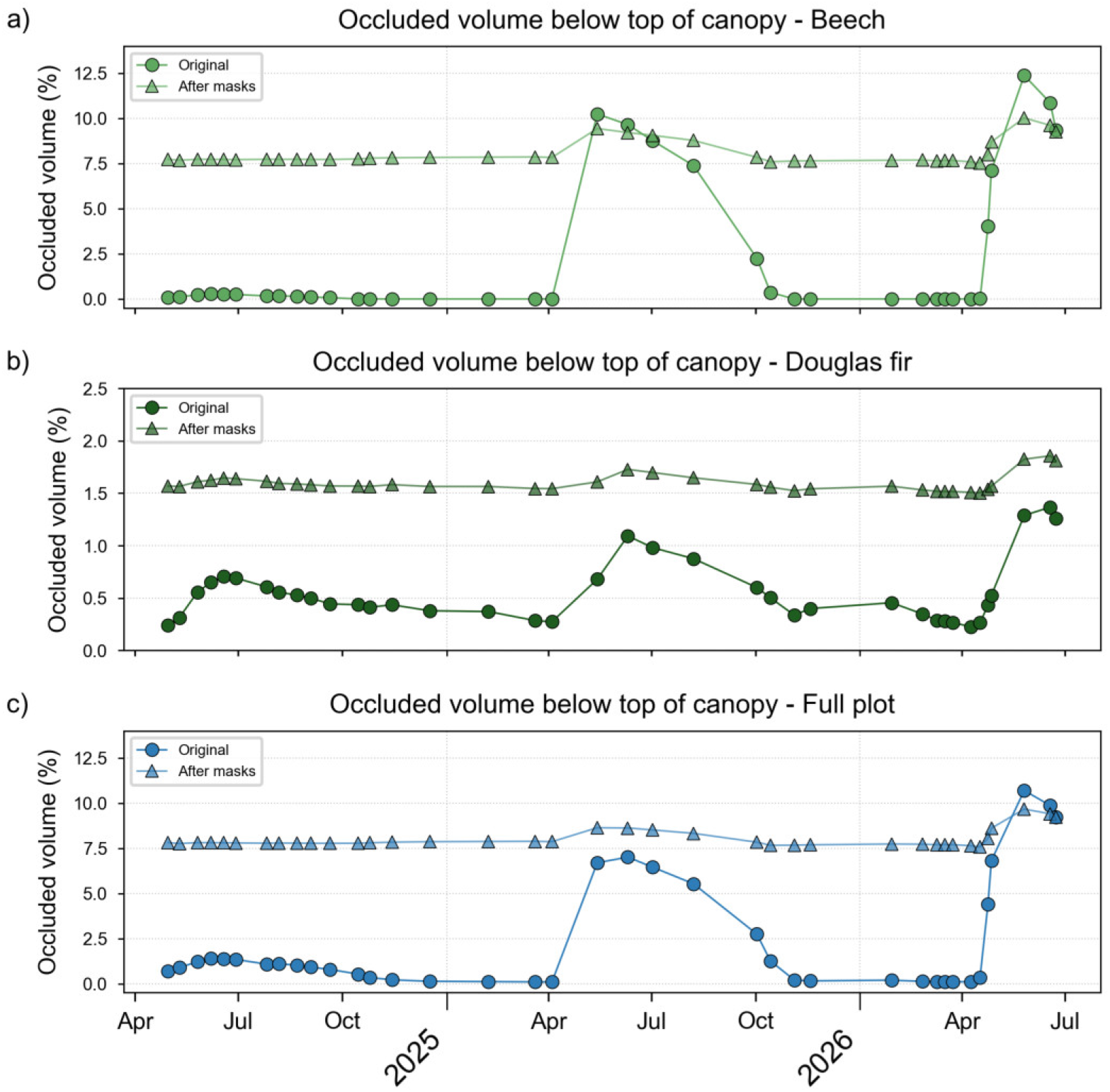
Percentage of volume occluded below the top of canopy in the a) beech patch, b) Douglas fir patch, and c) full plot. Lines with circular markers represent the occluded volume inherent to the original datasets, lines with triangular markers the occluded volume after applying the consistency mask and initializing empty space.

The full plot in Fig. 5c) displays a mixed influence from both tree species. The steady increase of the occlusion in the original datasets within each following summer is a further indicator of the increasing vegetation density. After applying the consistency masks for inter-flight comparability, the occluded volume remained mostly around 7.8 % across the time series, with slight summer maxima of up to 8.6 % in 2025 and 9.7 % in 2026.

### 3.2. Variations in vertical height profiles

The vertical distribution of forest density was analyzed by aggregating metrics along horizontal voxel layers. To fully illustrate the temporal dynamics of the vertical height profiles across the entire time series, animated videos are provided in Figs. 7 and 9. For a concise summary of the seasonal variability, the following figures illustrate the range of variance by displaying specific minima and maxima.

#### 3.2.1. Mean voxel transmittance per height

The mean voxel transmittance in Fig. 6 is aggregated along the horizontal layers within each AOI. As seen in subsubsection 3.1.1, the beech patch shows pronounced phenological shifts. The center of the canopy layer is located at approximately 26 m height, where transmittance is around 0.87 in winter and shifts from 0.55 in summer 2024 to 0.48 in summer 2025 and 0.5 in summer 2026. Comparing summer profiles, a strong decrease in transmittance is observed from 2024 to 2025, with a vertical upward shift of the transmittance distribution. This suggests an increase in total blocking biomass and overall tree growth. In the following summer 2026, transmittance reaches a similar minimum transmittance, yet, continues with an upward shift of the distribution. The comparison of winter profiles 2024/25 and 2025/26 also reflects this height shift. Transmittance below the canopy layer is close to one, indicating mainly permeable empty space.

**Figure 6.**
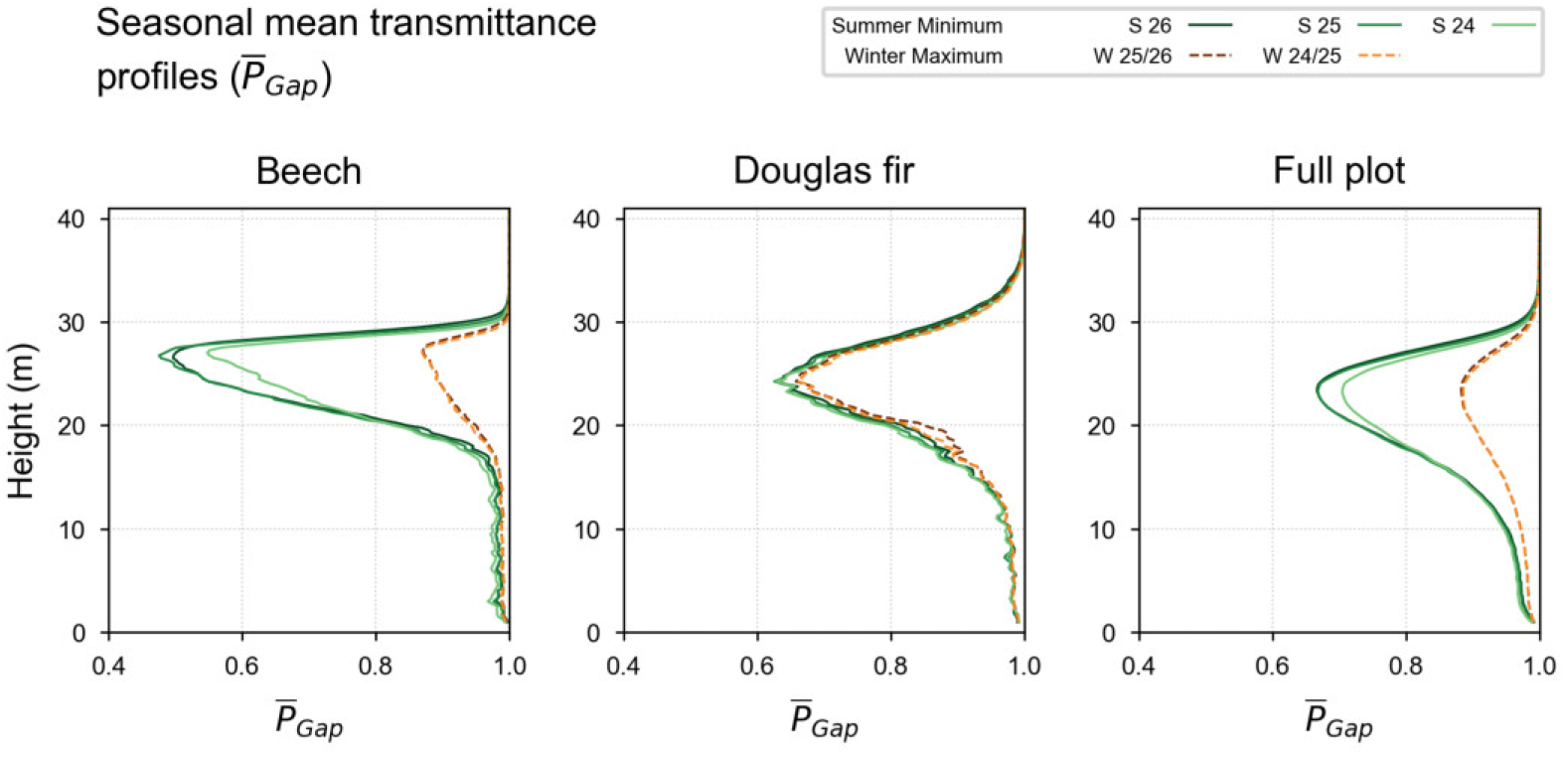
Mean transmittance height profiles with seasonal summer minima and winter maxima for the three AOIs.

The Douglas fir patch exhibits only subtle phenological changes, with a difference of ∼ 3 % between summer and winter transmittance. However, a similar vertical upward shift of the transmittance distribution is observed, which also indicates a general increase in tree height.

Again, the vertical profiles of the full canopy closely mirror those of the beech patch and appear smoother due to the averaging over a larger area. Transmittance minima are ∼ 0.88 in winter and ∼ 0.67 in summer 2025 and 2026. The distribution covering mid- and understory likely reflects mixing of species with varying heights in the full plot.

The full dynamics of *P_Gap_* height profiles of all acquisition dates can be observed in the the video provided in Fig. 7. During spring and autumn, height profile show rapid transitions, especially in AOIs with deciduous trees. During winter and summer, height profiles remain relatively stable, outlining the robustness of transmittance estimations.

**Figure 7.**
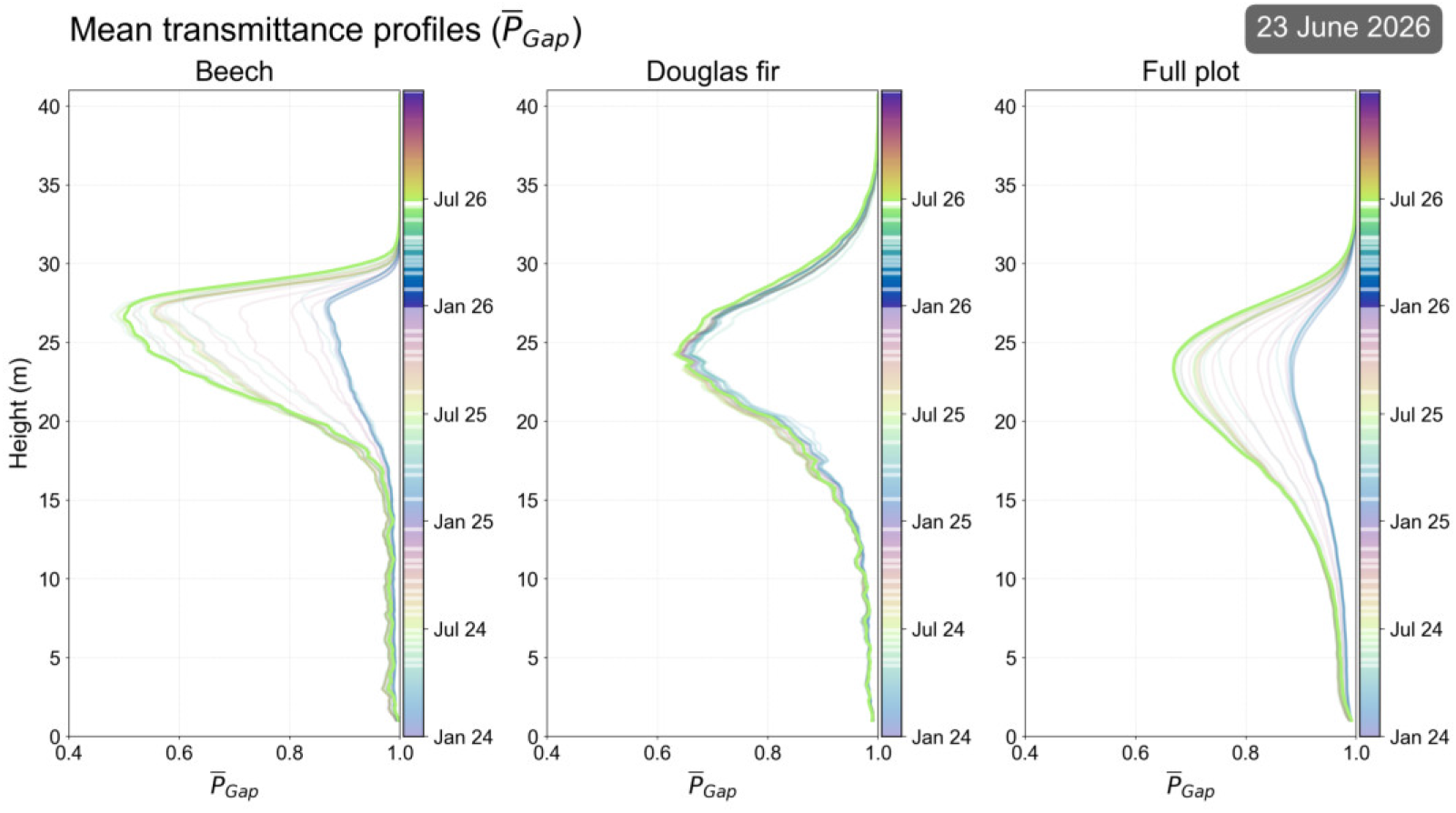
Video of consecutively following mean transmittance height profiles of all data acquisitions showing seasonal and across year variations.

[VIDEO]

#### 3.2.2. Cumulative attenuation volume per height

The vertical distribution of the cumulative attenuation volume (Σ*AV*) as a proxy for blocking biomass is shown in Fig. 8. The beech patch reveals a primary canopy layer centered at 27 m, where seasonal variations are strongest. An increase in attenuation is observed between summer 2024 and summer 2026, with a smaller increase in summer 2026. The upward shift of the Σ*AV* distribution is also visible, which aligns with the mean transmittance profiles (see Fig. 6). Below the main canopy layer, variations are small as most of biomass are likely woody elements not affected by phenology.

**Figure 8.**
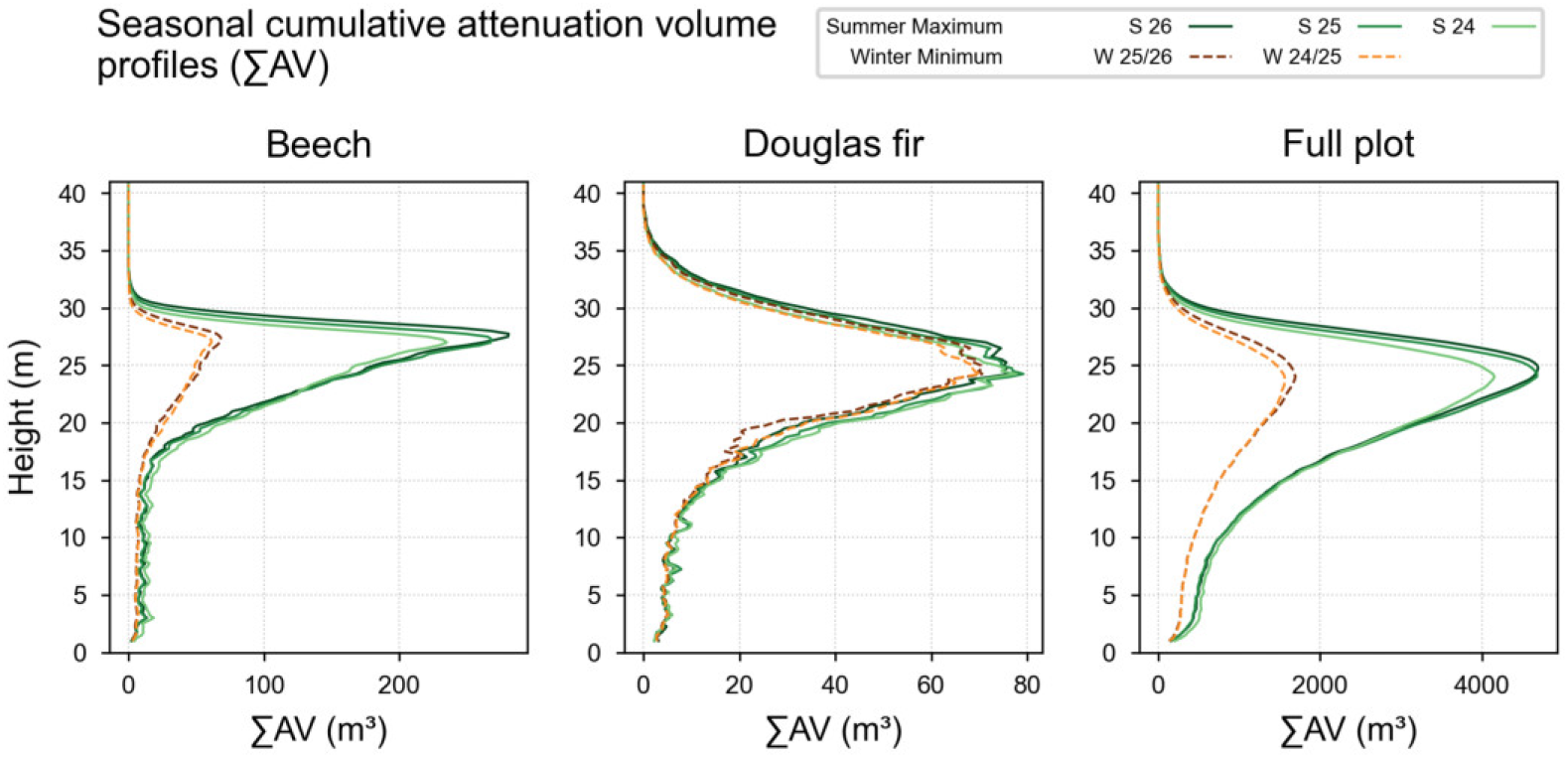
Cumulative attenuation volume height profiles with seasonal summer maxima and winter minima for the three AOIs.

The Douglas fir patch shows a much subtler phenological cycle, though a general increase of attenuation volume and a vertical upward shift of the distribution is visible. The full plot is again similar to the beech patch, with an overall increase in Σ*AV* and a vertical upward shift of the distribution. However, mid- and understory show again larger contributions to blocking biomass compared to the beech patch.

The full dynamics of Σ*AV* height profiles of all acquisition dates are illustrated in the video provided in Fig. 9. During spring, the rapid increase in canopy attenuation and filling from the higher layers of canopy is visible in the beech patch and full plot. Again, height profiles remain relatively stable during winter and summer, underlining the robustness of attenuation estimations.

**Figure 9:**
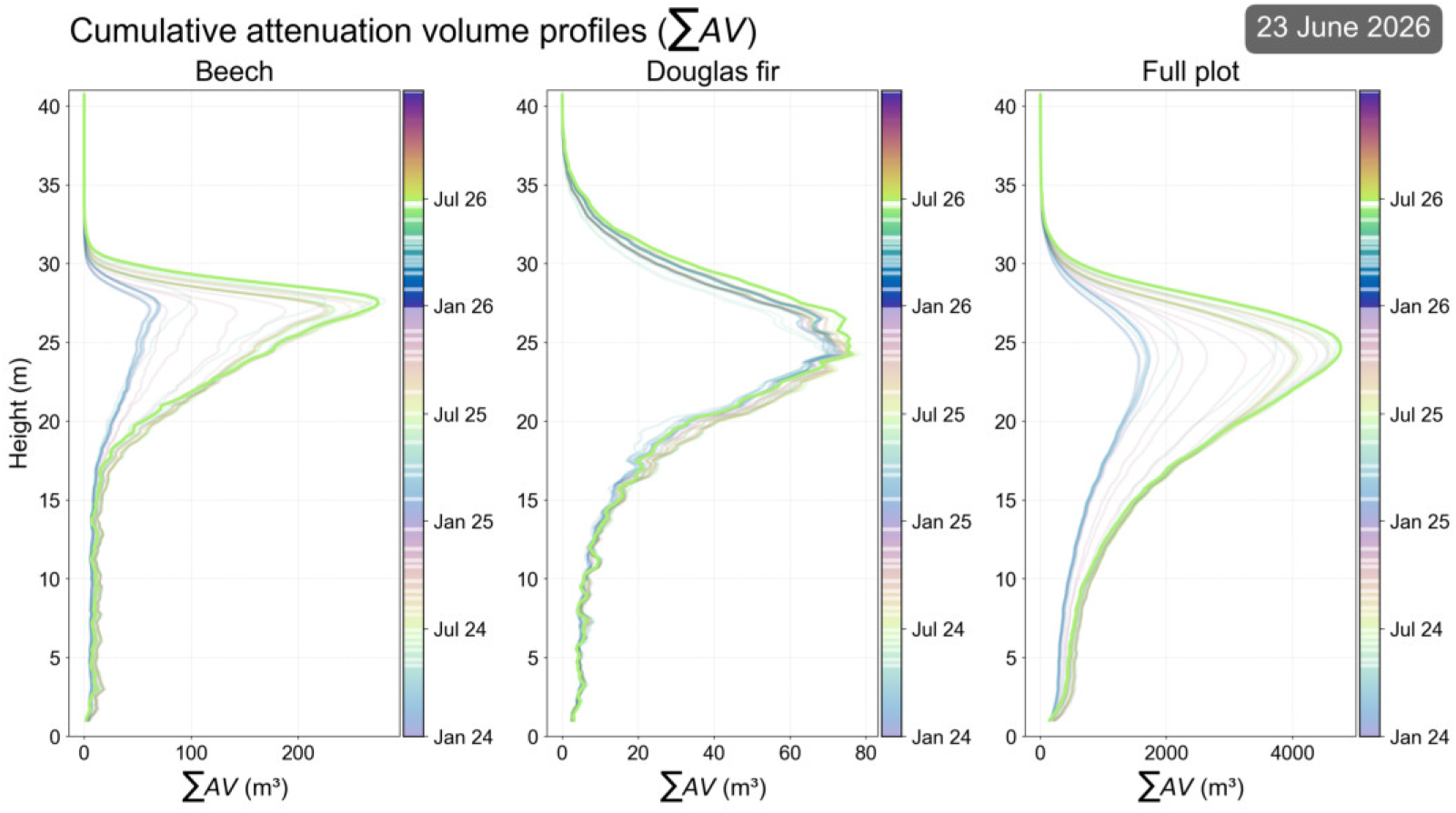
Video of consecutively following cumulative attenuation volume height profiles of all data acquisitions.

[VIDEO]

### 3.3. 2D spatio-temporal distribution of forest density

To analyze the horizontal heterogeneity of forest structure, we projected the 3D voxel space into high-resolution 2D maps by aggregating different metrics along vertical voxel columns.

#### 3.3.1. Cumulative Attenuation Volume Mapping

Summing the attenuation volume (Σ*AV*) along the vertical axis results in maps that represent the total light-blocking capacity of the vegetation. As established in subsection 2.6, the values of attenuation are used primarily for relative comparison of spatial distribution within a single date map and across the time series.

Two exemplary datasets, one from winter (2 February 2025) and one from summer (7 July 2025), are shown in Fig. 10. To enhance visualization and minimize the influence of outliers, the range is clipped to the 1st and 99th percentiles. The cumulative maps reveal a strong contrast between the deciduous and coniferous trees. During the winter (leaf-off) season, high attenuation of the evergreen species highlights the north-south row of Douglas fir in the center and the patch of Douglas fir and silver fir in the south-east of the plot. Their blocking capacity largely remains the same in summer and thus independent of the season.

**Figure 10.**
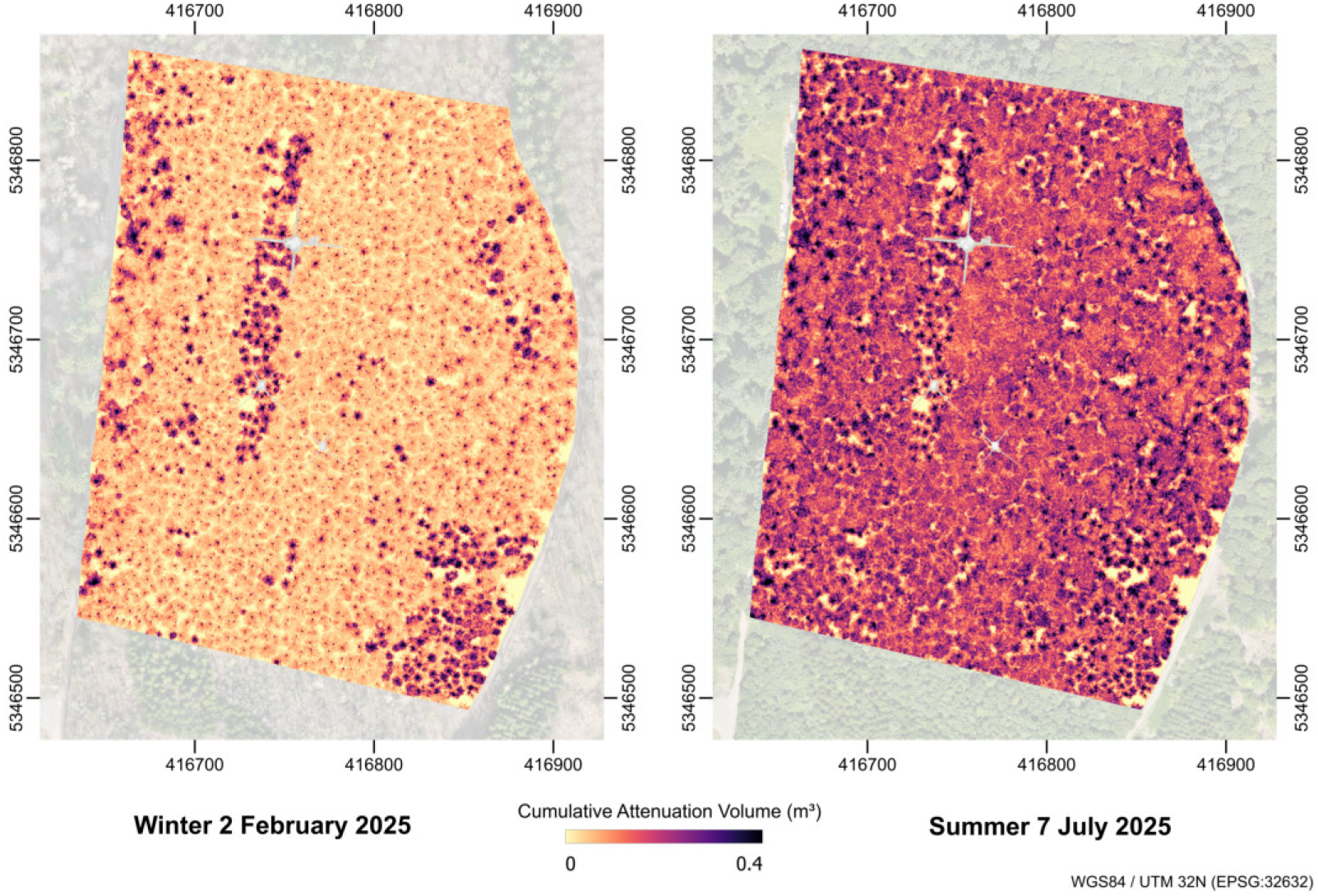
Maps with the cumulative attenuation volume aggregated along vertical voxel columns in 0.25 m resolution. The left map shows an exemplary winter acquisition from 2 February 2025. The right map shows an exemplary summer acquisition from 7 July 2025. The value range is clipped approximately to the 1st and 99th percentiles.

Fig. 10 shows a strong increase in attenuation in summer across the deciduous trees, where foliation closes previous gaps and creates a much denser canopy. However, some canopy gaps remain visible, allowing incident light to reach the forest floor.

The dynamics of individual Σ*AV* maps can be observed in the video provided in Fig. 11. Beyond the primary phenological cycle, the maps allow for the identification of tree-specific dynamics, such as an inhomogeneous start of leafing-out and leaf-fall across different individuals. Furthermore, the 2D distribution enables the detection of structural disturbances, such as the falling or removal of individual trees, which appear as sudden localized decreases in cumulative attenuation as indicated by arrows in Fig. 12.

**Figure 11.**
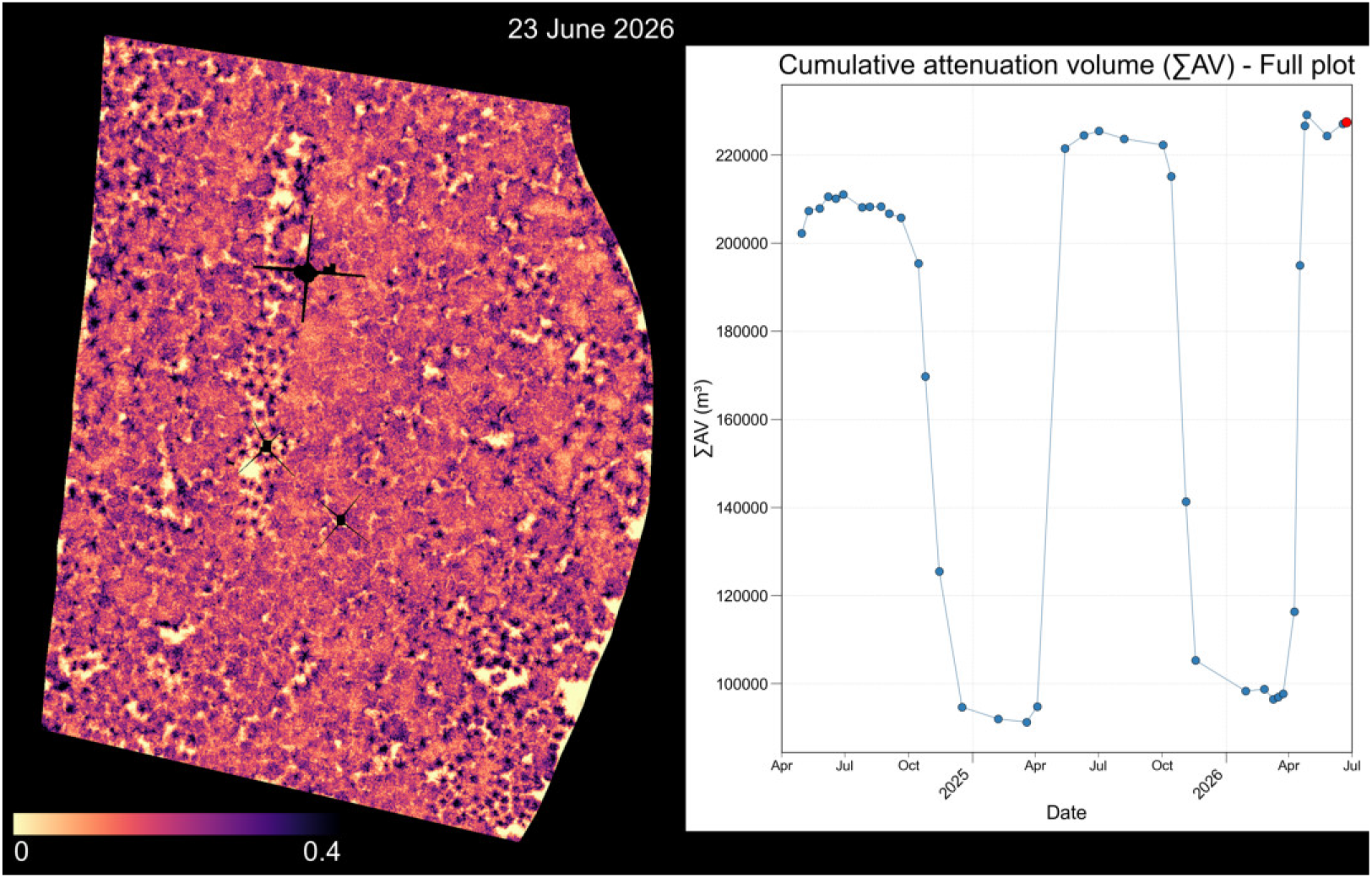
Maps with the cumulative attenuation volume aggregated along vertical voxel columns in 0.25 m resolution. The left panel shows the individual Σ*AV* map for each acquisition date. The value range is clipped approximately to the 1st and 99th percentiles. Throughout the time series, phenological change and distinct disturbance events are visible. The right panel shows the Σ*AV* for the full plot as presented in Fig. 4c).

**Figure 12.**
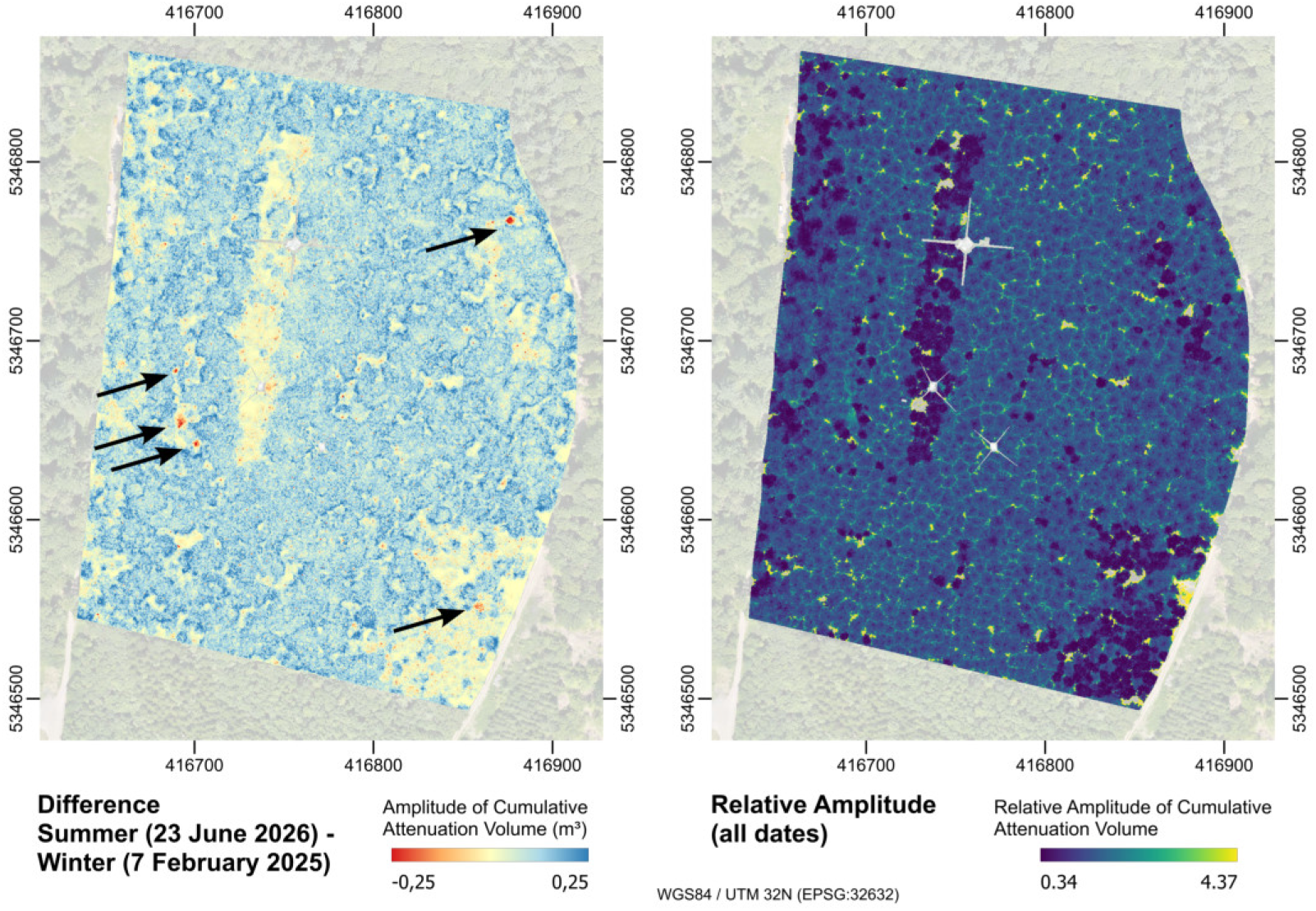
Left side: Exemplary map with a bi-temporal difference of the cumulative attenuation volume aggregated along vertical voxel columns. The maximum range was clipped to the ∼ 98th percentile and the minimum range value was adjusted to ensure proper centering of the diverging scale. Disturbances are highlighted by arrows. Right side: Relative amplitude of the cumulative attenuation volume (*RAV*) across all dates. Value range is clipped approximately to the 1st and 99th percentiles.

[VIDEO]

#### 3.3.2. Change in Cumulative Attenuation Volume

To further quantify the horizontal distribution of structural dynamics, we analyzed the seasonal change in cumulative attenuation volume (Σ*AV*) with an exemplary bi-temporal difference map and a relative amplitude map of the full time series.

The difference map Fig. 12 illustrates the change in Σ*AV* between a summer (23 June 2026) and a winter (7 February 2025) acquisition. To enhance the visualization of structural shifts and minimize the influence of outliers, the maximum range was clipped to the ∼ 98th percentile, with the minimum range value adjusted to ensure appropriate centering of the diverging scale. Deciduous trees show a strong increase in Σ*AV* due to their additional leaves, which is particularly pronounced at crown edges where foliage grew into gaps. In contrast, evergreen coniferous trees show little change in cumulative attenuation volume. Beyond phenology, the difference map highlights structural disturbances, which became already apparent in the animated video (Fig. 11). The falling of individual trees is clearly visible as localized losses in attenuation (highlighted by arrows), confirming the observations from the time-series animation.

Many tree stems in the difference map are characterized by distinct negative values. Because stems are not subject to phenological change, it indicates the presence of sampling and methodological artifacts as discussed later on. The map on the right in Fig. 12 shows the relative amplitude of the cumulative attenuation volume (*RAV*) across the full time series, visualized between the 1st and 99th percentile. While the difference map highlights the volume change, the *RAV* shows the phenological signal isolated from the absolute magnitude and highlights species-specific behavior. Coniferous trees are characterized by very low relative change, whereas deciduous trees show the highest *RAV* values, particularly in areas of crown expansion. This normalization effectively separates the volume impacted by seasonal phenology from the woody components of the deciduous trees, revealing the underlying tree architecture.

## 4. Discussion

This study provides a high-resolution perspective on the spatio-temporal dynamics of forest canopy transmittance across a two-year period with an extensive, consistent multi-temporal UAV LiDAR dataset. By maintaining consistent data acquisition and conducting an occlusion-aware analysis, this work demonstrates the potential of using relative transmittance as a direct proxy to quantify structural change. In the following sections, we discuss our results, methodological challenges, limitations, and the transferability of our work.

### 4.1. Spatio-temporal structural dynamics and phenological signatures

By combining high-resolution 3D voxel mapping with dense temporal sampling, this study enables a detailed interpretation of distinct phenological signatures and structural growth patterns, marking a transition toward the “fourth dimension” of LiDAR datasets in forestry (Calders et al., 2015).

The high-frequency sampling allowed the separation between seasonal phenological cycles and genuine structural growth. Both could be observed in the mean attenuation of filled voxels (*A̅_filled_*) as well as the cumulative attenuation volume (Σ*AV*), with a distinct contrast between deciduous and evergreen species. These results confirm that UAV LiDAR derived transmittance acts as a sensitive proxy of structural change.

The analysis of the beech patch showed a high-amplitude seasonal change in forest density. The rapid increase of *A̅_filled_* and Σ*AV* during spring phenology observed in our results aligns with Calders et al. (2015) where leafing of deciduous trees followed a sigmoidal curve. The rapid foliation changes observed between April and May highlight the necessity of adapting highfrequency acquisitions to the speed of phenological processes and structural change (Calders et al., 2015; Campos et al., 2021).

The observed vertical phenological variation in spring 2026, where canopy filing begins at the top and descends to the lower layers, aligns with Calders et al. (2015) who noticed that upper canopy layers have an earlier phenological onset than the understory. While Gressler et al. (2015) state that upper canopy layers also start earlier with autumn phenology, this was not observed by inspection of the height profile animation at the plot level (see Fig. 9). For this case, higher acquisition frequency and individual tree-level analysis might be necessary, as stand-level aggregates metrics may obscure these vertical nuances in autumn.

The increase in summer peak attenuation (*A̅_filled_* and Σ*AV*) from 2024 to 2025 indicates an overall increase in forest density beyond seasonal fluctuations. The gradual decline in attenuation during summer months may be attributed to environmental stressors or biomass loss, while the rapid autumn senescence reflects leaf fall driven by temperature, humidity and light availability (Delpierre et al., 2009; Gressler et al., 2015; Schnelle, 1955).

In contrast, the Douglas fir patch exhibited significantly lower variance and lacked a stable winter baseline. The later peak attenuation in Douglas fir (June) compared to beech (May) supports findings of delayed growth onset in conifers relative to deciduous species (Bärwald et al., 2026). While Gould et al. (2012) note that basal-area growth periods can extend beyond standard definitions (e.g. from late February until November), the variation in LiDAR derived voxel transmittance is likely driven by foliage and twig dynamics rather than basal growth.

The delayed and longer growth period results from biological differences between species, which could reflect a strategy of complementary resource use in time to reduce direct competition for light and water during peak biological activity (Bärwald et al., 2026). Moreover, observed phenological signatures may not be universal but dependent on management history and silviculture, which can alter senescence phases and growth patterns (Bajocco et al., 2026).

The data reveals clear signatures of inter-annual structural growth. The consistent increase in peak attenuation from 2024 to 2026, coupled with the upward vertical shift in transmittance profiles, indicates an increase in blocking biomass and total tree height.

The sudden increase in *A̅_filled_* and Σ*AV* as well as occlusion in the Douglas fir patch during early 2026 indicate structural change that affected transmittance estimations. Since this signal is missing in the beech patch and is found in the 1-2 preceding and consecutive data acquisitions, it is likely not a sampling error. Literature on species cambial and foliar phenology suggest that growth initiation is strictly governed by chilling and heat-forcing requirements (Campbell, 1974; Campbell and Sugano, 1975) with species specific temperature threshold (e.g., accumulation of ∼ 1200 h at 0–5^◦^ C (Bailey and Harrington, 2006)). In January, those chilling requirements were not satisfied, wherefore these models could not explain a sudden growth of Douglas fir. Hence, this phenomenon will be subject of future research linking structural LiDAR data with eco-physiological data collected via the in-situ sensor network at the field site, potentially laying the foundation for the development and validation of phenological growth models at scale.

The 2D mapping of Σ*AV* reveals significant intra-stand variability. For deciduous trees, an inhomogeneous start of leafing-out and senescence can be observed across individual trees Fig. 9. This aligns with Malyshev et al. (2022), who demonstrated significant inter-individual bud burst variation of up to three weeks within a single stand, attributed to genetic differences and warming rates.

The high-resolution ULS data enables the detection of discrete disturbance events distinct from phenological changes, such as tree fall and branch losses as voxel attenuation drops. This capability underscores the advantage of ULS to coarser satellite data to monitor small-scale structural dynamics (Calders et al., 2015).

The relative amplitude map aggregated across the dates effectively isolates the phenological signal (high for deciduous) from volume covered by mainly woody components, showing the ability to analyze species-specific dynamics without prior segmentation.

### 4.2. Occlusion and sampling bias

While the relative amplitude effectively captures phenological dynamics, the 2D difference map reveals a distinct methodological artifact regarding the sampling of woody biomass. Specifically, tree stems consistently exhibit lower attenuation in summer than in winter, appearing as prominent negative values in the summer–winter difference maps. Since woody stems do not undergo phenological change, this indicates a methodological sampling bias where foliage occlusion in summer prevents reliable stem sampling compared to winter leaf-off conditions.

The original occlusion increased from negligible levels in summer 2024 to over 12% in the beech patch by 2026 which highlights a fundamental challenge in long-term structural monitoring: Structural growth directly alters the volume of effectively sampled space. This creates a observation bias, where the most relevant forest volume with the strongest growth is also the most severely affected by increased occlusion, potentially masking key structural dynamics. While the applied consistency mask (excluding voxels occluded *>* 3 times) and initialization of empty space reduced occlusion variation (stabilizing around 8–10%), it excludes relevant structural information, highlighting the trade-off between comparability and data completeness. The application of a stricter occlusion mask (excluding voxels occluded ≥ 1 times) could reduce this bias, however inadvertently filters out even more areas subject to structural change and thus compromises the comparability of datasets through excessive data loss.

In consequence, these findings indicate an overall underestimation of attenuation due to occlusion throughout the study. This effect is for ULS particularly pronounced below dense canopy layers and for deciduous trees larger during summer months. Furthermore, while *A̅_filled_* decreases during summer, Σ*AV* remains on a plateau. This suggests that the reduction in mean attenuation may be compensated by improved beam penetration, which increases the volume of effectively sampled voxels and stabilizes the cumulative attenuation volume. It is highly probable, that similar sampling effects occur within the denser parts of the upper canopy, even though the bias is masked by the strong phenological change. However, this might explain the weak growth signal observed between summer 2025 and 2026, compared to the increase from 2024 to 2025. Despite little phenological change, a similar bias was observed in coniferous trees, suggesting that the surrounding deciduous foliage may obstruct laser penetration to conifer stems. Ultimately, winter “leaf-off” conditions likely provide the most reliable baseline for sampling stems and woody architecture.

This demonstrates that in dense, evolving forests, any multi-temporal analysis must account for the fact that occluded forest volume is systematically tied to the structural evolution of the canopy. Consequently, relying on simple point cloud statistics without an occlusion-aware framework risks misinterpreting a loss of observable volume as a loss of physical biomass.

### 4.3. Methodological robustness and transferability

A core contribution of this work is the deliberate analysis of relative transmittance change instead of assuming the modeling of absolute PAD or LAD values. While PAD is typically considered a meaningful ecological trait, its retrieval relies on assumptions regarding leaf angle distribution and clumping coefficients, which differ by species and even vary at the voxel level. Furthermore, Jensen’s inequality of the convex beer-lambert function leads to a systematic positive bias in PAD estimates due to sampling variance, an increased uncertainty in low transmittance voxels, and a voxel size dependent underestimation when averaging heterogeneous canopy elements (Arnqvist et al., 2020; Pimont et al., 2019; Verley, 2026).

This choice is further justified by the technical infeasibility of retrieving ground truth data in high-resolution voxel space. Destructive sampling is only conducted at aggregated scales (e.g. Béland et al., 2011) or on small samples. Because absolute values are sensitive to forest structure and sensor settings, method development and validation often lack transferability. Validation methods, such as LAI-2200 or DHP again provide only aggregated values on small scales and possess their own inherent flaws and biases, questioning their use for absolute calibration.

Consequently, using relative transmittance provides a more robust representation of structural shifts, as it tracks the LiDAR system’s actual penetration capabilities. The extensive time series of 38 flights acquired with identical sensors and harmonized settings establishes a robust baseline, where the consistency of the signal validates the observed dynamics even if absolute values remain uncalibrated.

The insights obtained on structural dynamics are substantial, yet remain subject to plot-specific constrains such as local forest structure and leaf angle distributions, both impacting the measured transmittance. Furthermore, the observed biases and occlusion patterns are partly tied to the technical specifications of the DJI Zenmuse L2 sensor, including its wavelength, footprint size, and viewpoint geometry.

Beyond these constraints, this study identified several persistent challenges inherent in any multi-temporal LiDAR analysis of forest density. For instance, LiDAR data acquisitions frequency must be enhanced to match the speed of structural change during phenological phases, yet sampling remains limited by a strict dependence on stable weather conditions. Moreover, the transition to newer sensor generations (e.g. DJI Zenmuse L3) often hinders long-term ecological monitoring by introducing inconsistencies in data acquisition. Finally, the analysis of high-resolution voxel space via direct voxel comparison remains restricted by subtle movements of forest structure, such as branch swaying or sagging due to increased leaf weight. Despite these methodological obstacles encountered, this study provides a valuable framework for the analysis of multi-temporal LiDAR data in complex forest environments.

Beyond the observed phenological dynamics and methodological challenges, this work demonstrates that with consistent data acquisition, relative transmittance patterns remain robust and can be transformed into a powerful tool for monitoring forest change over time. The use of high-frequency ULS acquisitions lays the foundation for explicit 3D mapping of forest transmittance with unprecedented spatial and temporal resolution. Furthermore, while the processing of high-resolution voxel space remains memory-intensive, the custom Python implementation developed for this study proves that such a workflow is scalable to the plot level. By navigating the trade-offs between sampling frequency and occlusion, this approach provides a pathway for implementing long-term, high-resolution structural monitoring across diverse forest ecosystems.

## 5. Conclusions and Outlook

This study investigated the spatio-temporal dynamics of forest canopy transmittance in a European mixed temperate forest using a consistently acquired extensive multi-temporal UAV LiDAR dataset comprising 38 flights over more than two years. With a custom Python implementation (CANOPy) we estimated voxel transmittance of the forest plot at 0.25 m resolution and performed occlusion classification and mapping.

We established an occlusion-aware framework, enabling robust multitemporal comparability while distinguishing true structural dynamics from sampling biases inherent in multi-temporal LiDAR datasets of forest environments. In this work we analyzed the relative change of voxel transmittance and voxel attenuation as a proxy for forest density by addressing three primary objectives:

First, we quantified temporal structural dynamics at the plot level and found distinct phenology signatures between deciduous and evergreen species. European beech exhibited high-amplitude seasonal dynamics with a 70% reduction in attenuation volume from summer to winter, whereas Douglas fir showed significantly lower variance (12.7%) and a delayed onset of growth. Furthermore, we detected an inter-annual increase of attenuation and inherent occlusion, both indicating an increase in blocking biomass.

Second, we analyzed variations in vertical height profiles and found a consistent upward shift of the transmittance distribution over the study period as well as an increase in maximum attenuation, underlining tree growth and crown expansion. Winter and summer baselines remained stable across years, validating the robustness of the transmittance estimation and the effectiveness of the occlusion-aware framework in maintaining comparability across acquisitions.

Third, we mapped the 2D spatial distribution, enabling a bi-temporal comparison to detect discrete structural disturbances, such as tree fall events. Moreover, cumulative attenuation maps highlight intra-stand variability, including individual tree-level differences in leafing-out and senescence timing. Relative amplitude maps effectively isolated the phenological signal from woody components, revealing species-specific behavior without prior segmentation.

This work demonstrates that high-frequency UAV LiDAR acquisitions can track forest structural dynamics with unprecedented spatio-temporal resolution. While absolute transmittance values may not be comparable across different sensor and acquisition parameters, the relative change is a powerful tool to understand structural forest dynamics. Furthermore we highlight that occluded forest volume is systematically tied to the structural evolution of the canopy and needs to be taken into account for multi-temporal LiDAR analysis.

In future studies, we aim to link the observed structural dynamics to ecophysiological measurements from the in-situ sensor network with parameters such as soil moisture, temperature and sap flow data. Examining the interrelationship between structural dynamics and environmental drivers will contribute to a more holistic understanding of forest ecosystems. Additionally, once automatic tree segmentation methods become sufficiently reliable, they will enable the investigation of forest dynamics at the individual tree level, unlocking the full potential of high-resolution capacity of LiDAR data. Finally, bridging satellite remote sensing observations with close-range ULS LiDAR and ground-based sensor networks has the potential to provide unprecedented insights into sustainable forest management and advance our fundamental understanding of ecosystem responses to environmental change.

## CRediT authorship contribution statement

Matthias Gassilloud: Conceptualization, Formal analysis, Investigation Data Collection, Methodology, Software, Validation, Visualization, Writing – original draft, Writing – review & editing.

Barbara Koch: Funding acquisition, Supervision, Writing – review & editing.

Anna Göritz: Conceptualization, Funding acquisition, Methodology, Project administration, Resources, Supervision, Validation, Visualization, Writing – original draft, Writing – review & editing.

## Acknowledgements

This work was funded by the Deutsche Forschungsgemeinschaft (DFG) Project-ID 459819582 SFB 1537. The authors would like to thank the “XR Future Forest Lab” funded by the Eva Mayr-Stihl Stiftung for the provision of the L2 sensor.

## Data availability

Both the data and the code are made publicly available. Data can be found [HERE link will be updated] and the code is published here: https://github.com/MGEOS/CANOPy.

## Appendix A. Example images of study site

**Figure A.13.**
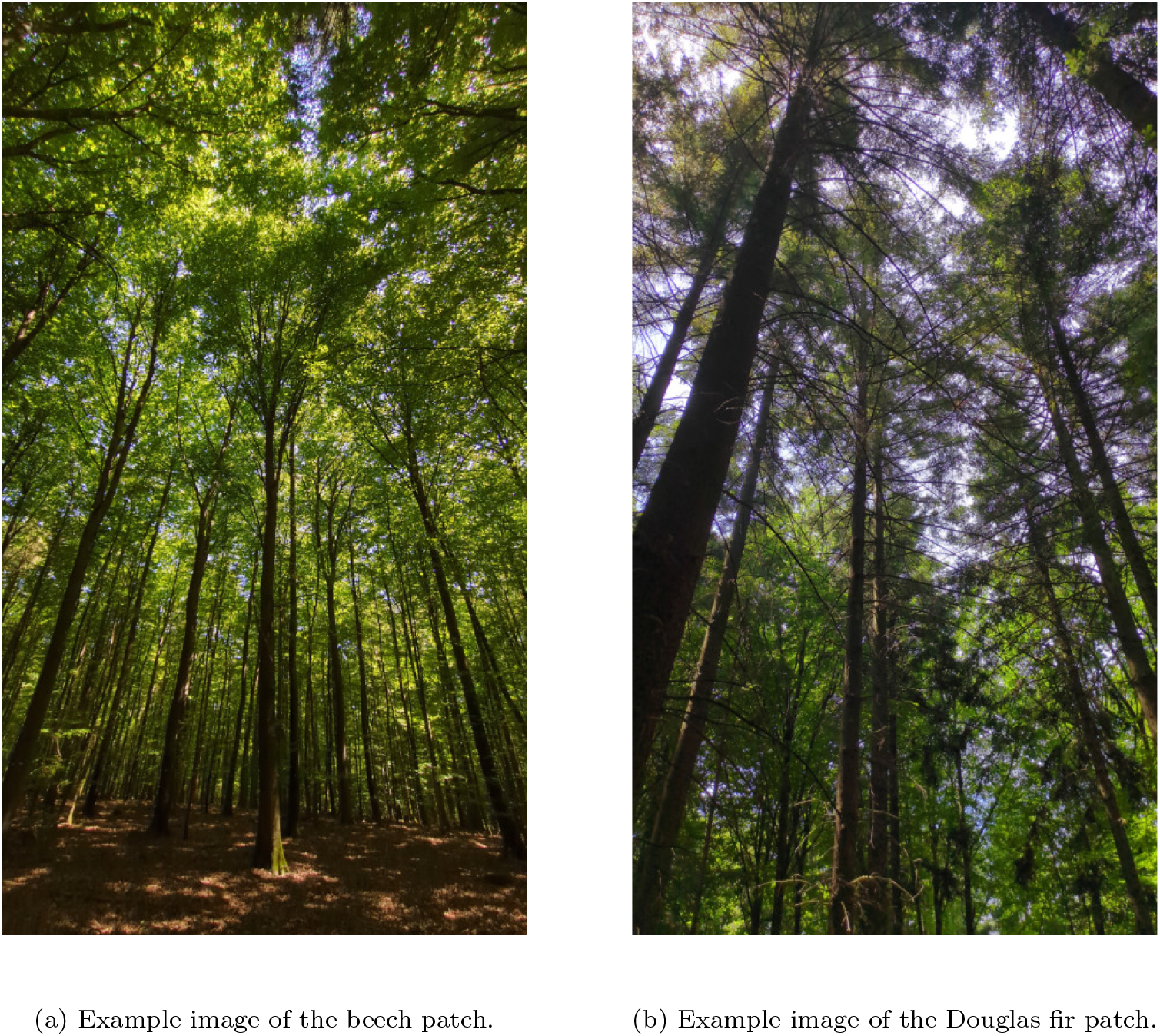
Example images of forest structure in the study site. The left image shows part of the beech patch. The right image shows part of the Douglas fir patch. Both patches have little mid- and understory.

## Appendix B. Overview of conducted flights

**Table B.2:**
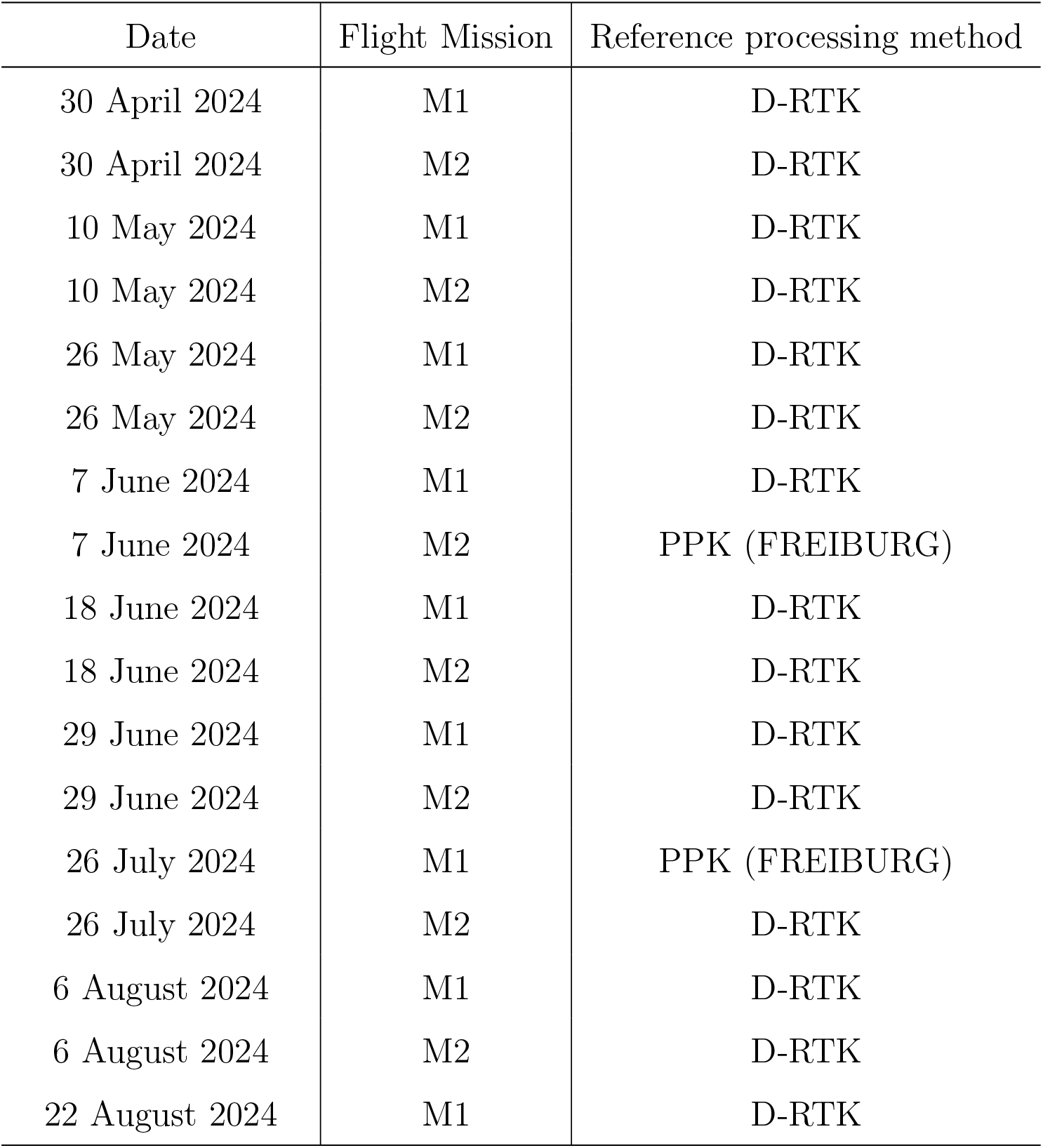
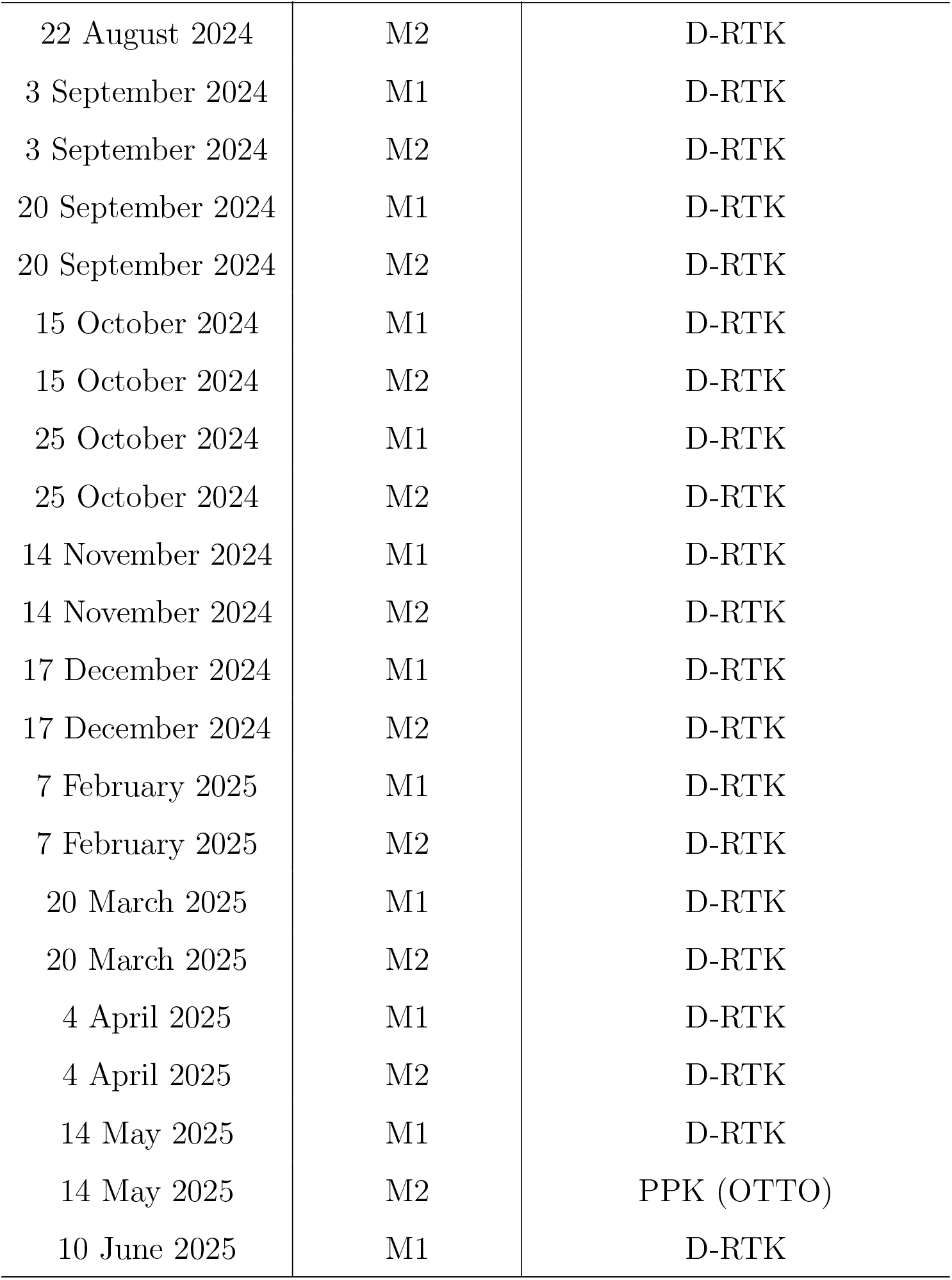
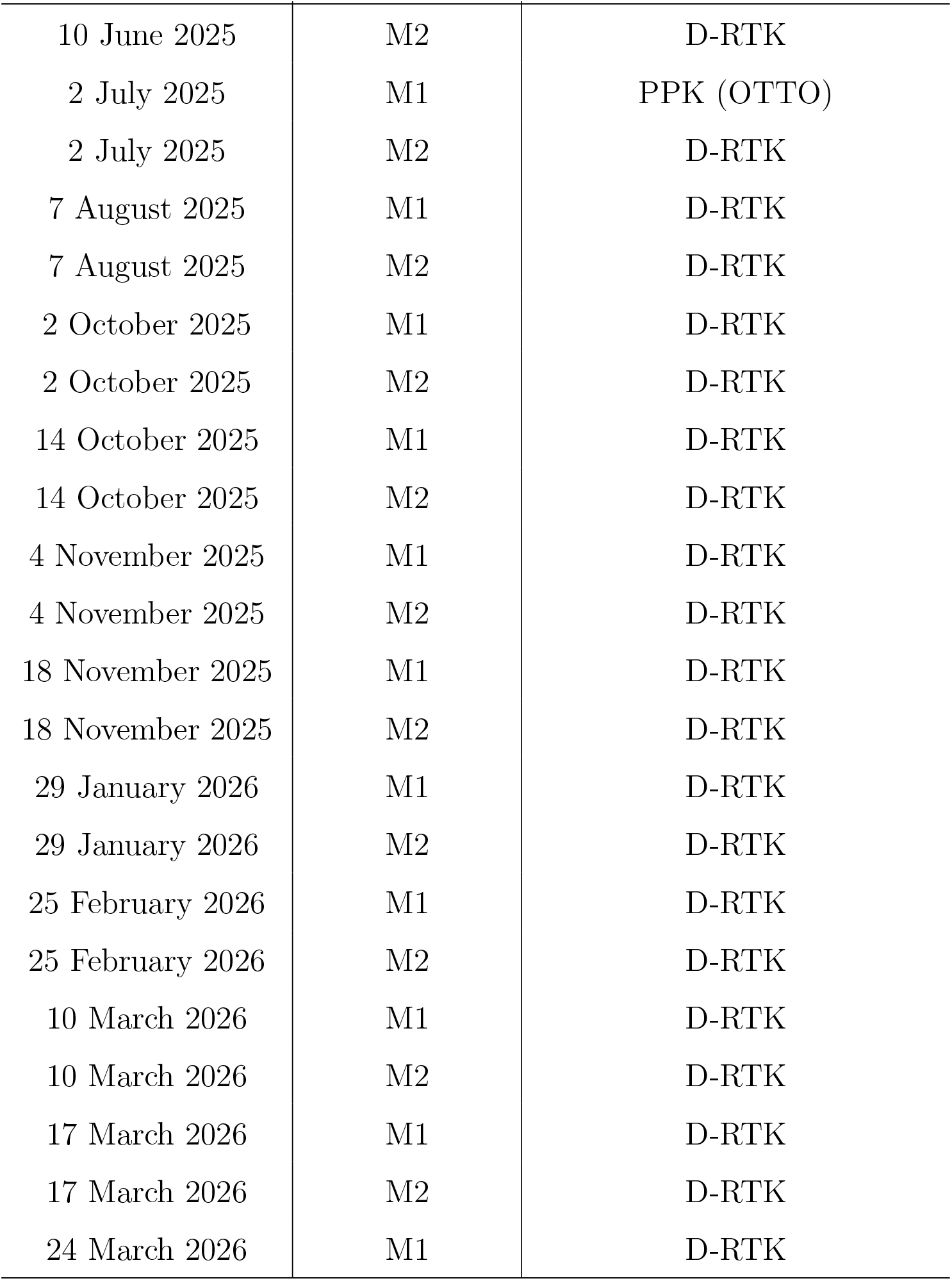
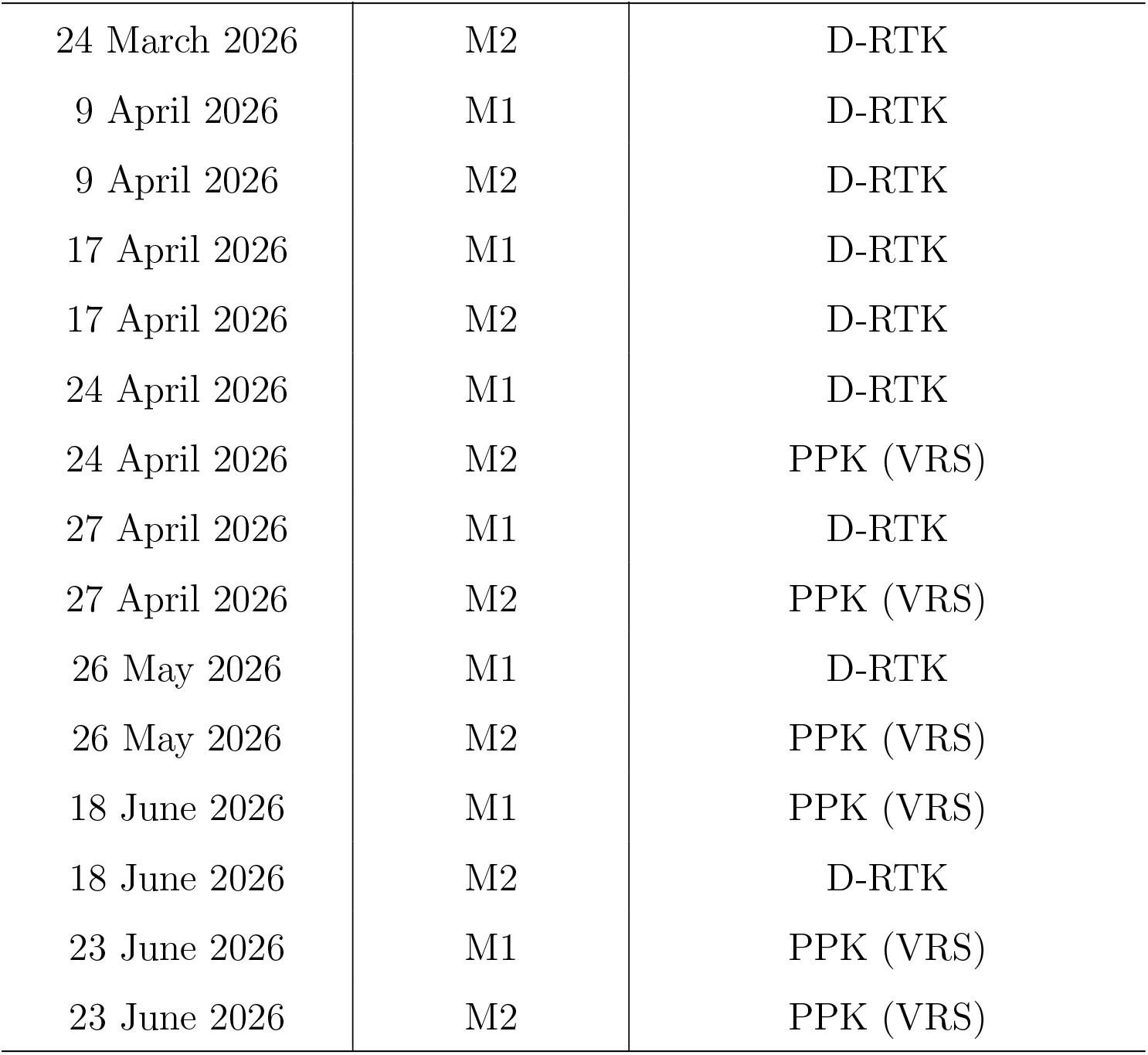
Overview of flight missions. All flights were conducted with D-RTK. However, in some cases point cloud reconstruction failed and had therefore to be processed with PPK using a (virtual) reference station. From 7th June 2024 until 9 August 2024, the FREIBURG SAPOS reference station was used. From 9 August 2024 until 31 December 2025 the OTTO SAPOS reference stations was used. From 2026 onward a virtual reference station (VRS) was calculated for the position where the D-RTK base stations has been placed.

| Date | Flight Mission | Reference processing method |
| --- | --- | --- |
| 30 April 2024 | M1 | D-RTK |
| 30 April 2024 | M2 | D-RTK |
| 10 May 2024 | M1 | D-RTK |
| 10 May 2024 | M2 | D-RTK |
| 26 May 2024 | M1 | D-RTK |
| 26 May 2024 | M2 | D-RTK |
| 7 June 2024 | M1 | D-RTK |
| 7 June 2024 | M2 | PPK (FREIBURG) |
| 18 June 2024 | M1 | D-RTK |
| 18 June 2024 | M2 | D-RTK |
| 29 June 2024 | M1 | D-RTK |
| 29 June 2024 | M2 | D-RTK |
| 26 July 2024 | M1 | PPK (FREIBURG) |
| 26 July 2024 | M2 | D-RTK |
| 6 August 2024 | M1 | D-RTK |
| 6 August 2024 | M2 | D-RTK |
| 22 August 2024 | M1 | D-RTK |
| 22 August 2024 | M2 | D-RTK |
| 3 September 2024 | M1 | D-RTK |
| 3 September 2024 | M2 | D-RTK |
| 20 September 2024 | M1 | D-RTK |
| 20 September 2024 | M2 | D-RTK |
| 15 October 2024 | M1 | D-RTK |
| 15 October 2024 | M2 | D-RTK |
| 25 October 2024 | M1 | D-RTK |
| 25 October 2024 | M2 | D-RTK |
| 14 November 2024 | M1 | D-RTK |
| 14 November 2024 | M2 | D-RTK |
| 17 December 2024 | M1 | D-RTK |
| 17 December 2024 | M2 | D-RTK |
| 7 February 2025 | M1 | D-RTK |
| 7 February 2025 | M2 | D-RTK |
| 20 March 2025 | M1 | D-RTK |
| 20 March 2025 | M2 | D-RTK |
| 4 April 2025 | M1 | D-RTK |
| 4 April 2025 | M2 | D-RTK |
| 14 May 2025 | M1 | D-RTK |
| 14 May 2025 | M2 | PPK (OTTO) |
| 10 June 2025 | M1 | D-RTK |
| 10 June 2025 | M2 | D-RTK |
| 2 July 2025 | M1 | PPK (OTTO) |
| 2 July 2025 | M2 | D-RTK |
| 7 August 2025 | M1 | D-RTK |
| 7 August 2025 | M2 | D-RTK |
| 2 October 2025 | M1 | D-RTK |
| 2 October 2025 | M2 | D-RTK |
| 14 October 2025 | M1 | D-RTK |
| 14 October 2025 | M2 | D-RTK |
| 4 November 2025 | M1 | D-RTK |
| 4 November 2025 | M2 | D-RTK |
| 18 November 2025 | M1 | D-RTK |
| 18 November 2025 | M2 | D-RTK |
| 29 January 2026 | M1 | D-RTK |
| 29 January 2026 | M2 | D-RTK |
| 25 February 2026 | M1 | D-RTK |
| 25 February 2026 | M2 | D-RTK |
| 10 March 2026 | M1 | D-RTK |
| 10 March 2026 | M2 | D-RTK |
| 17 March 2026 | M1 | D-RTK |
| 17 March 2026 | M2 | D-RTK |
| 24 March 2026 | M1 | D-RTK |
| 24 March 2026 | M2 | D-RTK |
| 9 April 2026 | M1 | D-RTK |
| 9 April 2026 | M2 | D-RTK |
| 17 April 2026 | M1 | D-RTK |
| 17 April 2026 | M2 | D-RTK |
| 24 April 2026 | M1 | D-RTK |
| 24 April 2026 | M2 | PPK (VRS) |
| 27 April 2026 | M1 | D-RTK |
| 27 April 2026 | M2 | PPK (VRS) |
| 26 May 2026 | M1 | D-RTK |
| 26 May 2026 | M2 | PPK (VRS) |
| 18 June 2026 | M1 | PPK (VRS) |
| 18 June 2026 | M2 | D-RTK |
| 23 June 2026 | M1 | PPK (VRS) |
| 23 June 2026 | M2 | PPK (VRS) |

## Appendix C. Voxel occlusion

**Figure C.14:**
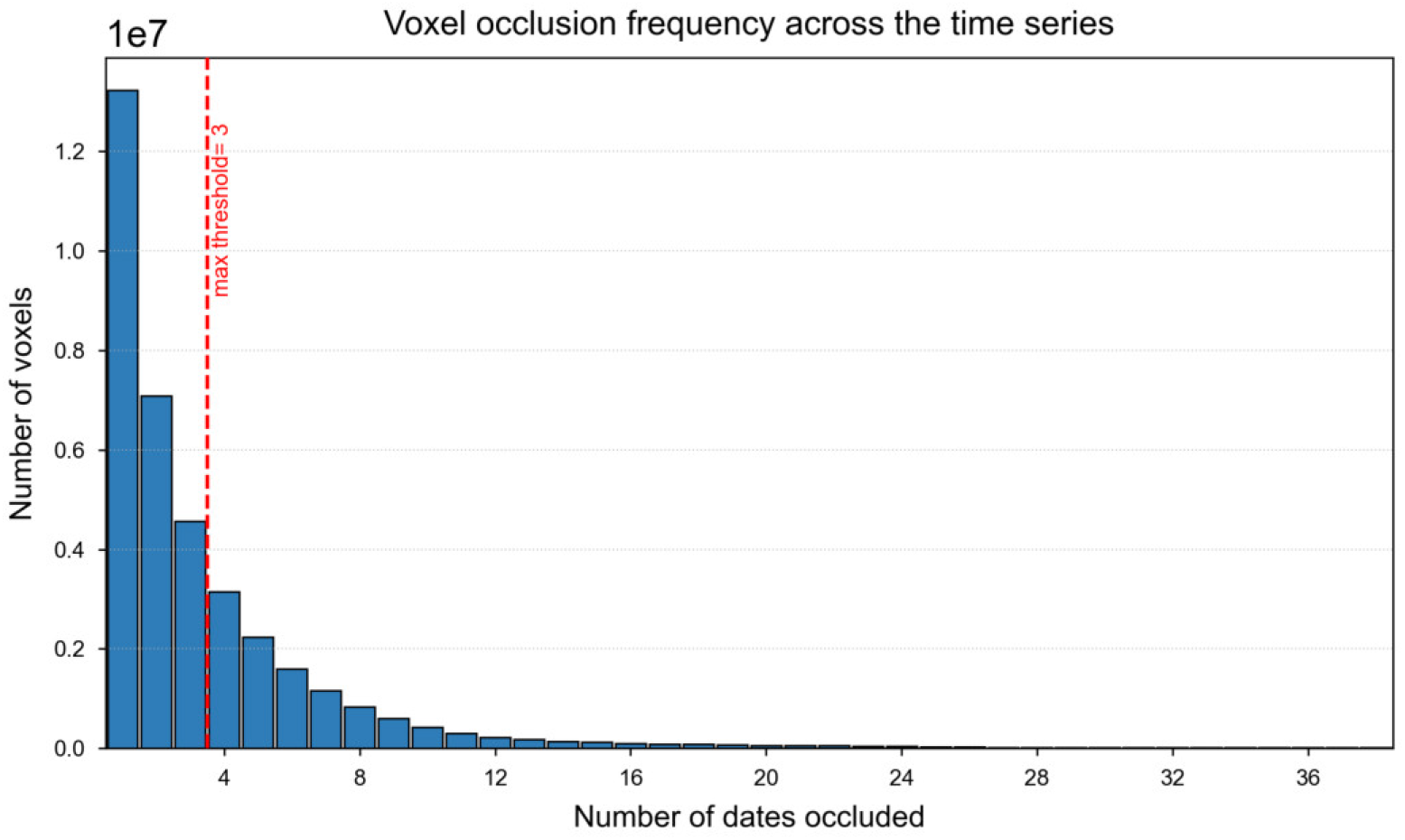
The frequency of voxel occlusion across the time series. Most voxels are occluded ≤ 3 times and were considered for analysis, while those occluded > 3 times were excluded (see subsection 2.5).

**Figure C.15:**
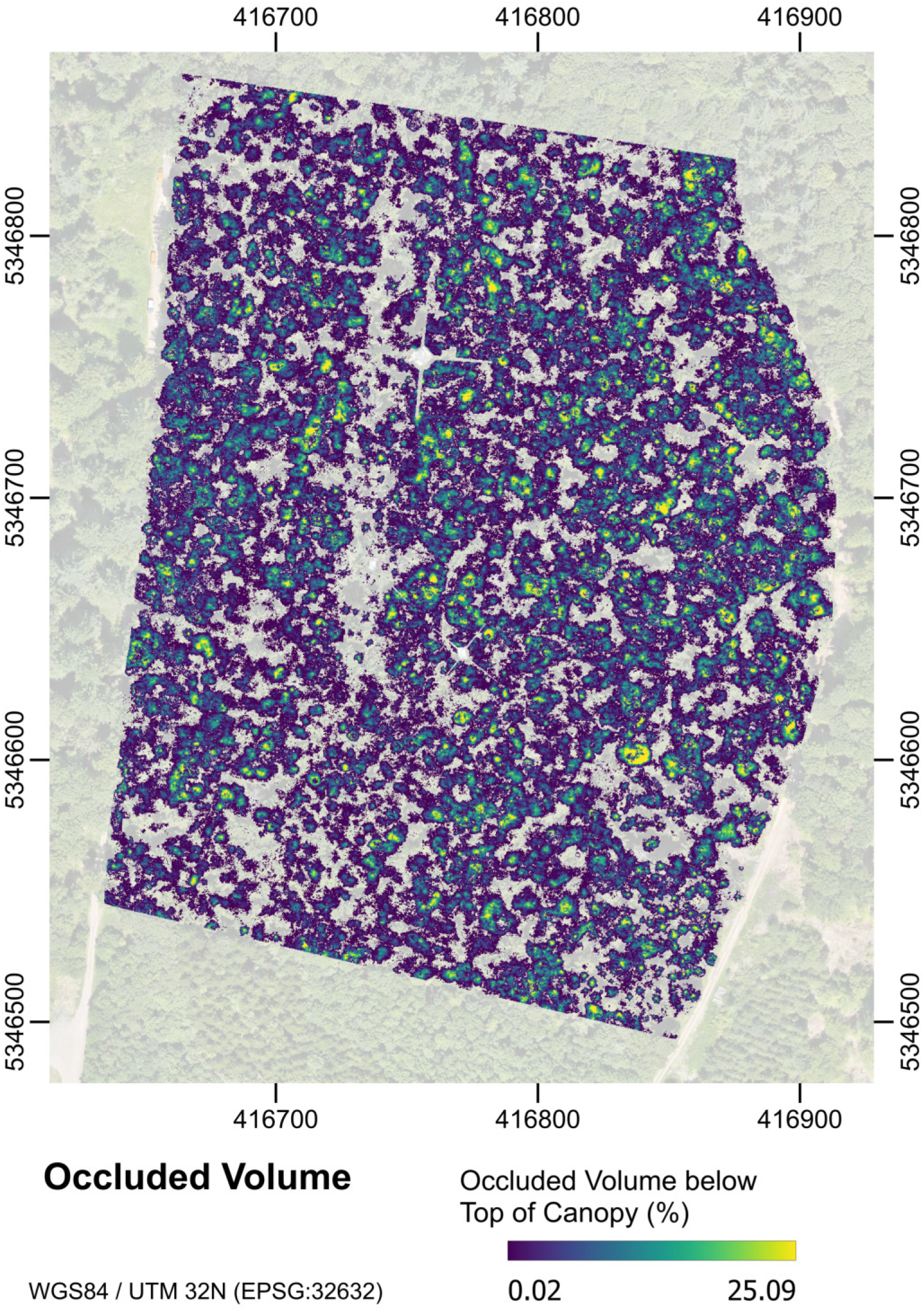
Mean occluded volume below top of canopy across all dates after application of exclusion rules (see subsection 2.5). Color scaling is clipped to the 1st to 99th percentile for visualization.

